# Anti-Z-NA antibodies can distinguish accessible Z-form states across DNA, RNA, and DNA–RNA substrates

**DOI:** 10.64898/2026.07.22.740205

**Authors:** Danielle Chin, Yongbo Luo, Zengyting He, Chaoran Yin, Riley M. Williams, Peter Dröge, Siddharth Balachandran, Dahai Luo

**Affiliations:** Lee Kong Chian School of Medicine, Nanyang Technological University, 59 Nanyang Drive, Singapore 636921; Institute of Structural Biology, Nanyang Technological University, 59 Nanyang Drive, Singapore 636921; Center for Immunology, Fox Chase Cancer Center, Philadelphia, PA 19111, United States; LambdaGentherapeutics Pte. Ltd., 1557 Keppel Road, Singapore 089066; National Centre for Infectious Diseases, 16 Jln Tan Tock Seng, Singapore 308442

**Keywords:** Z-RNA, Z-DNA, Z-DNA-RNA hybrids, anti Z-NA antibodies, ZBP1, ADAR1, Necroptosis

## Abstract

Left-handed (Z-form) double-stranded nucleic acid conformers (Z-DNA and Z-RNA, collectively called Z-NA) are molecular patterns recognized by ADAR1 and ZBP1, which are sensor proteins involved in innate immunity. Monoclonal antibodies raised against Z-DNA, such as Z22 and Z-D11, are employed as probes for the study of Z-NAs, but their substrate specificity and functional equivalence remain unclear. Here, we used biochemical, biophysical, and structural approaches to compare the binding modes and target specificies of Z22 and Z-D11, using Z-prone CG-rich dsDNA, dsRNA, and DNA–RNA hybrids as substrates. Both antibodies failed to form stable complexes with short CG-rich dsRNA under conditions compatible with antibody stability, which suggests that they are unable to induce A-to-Z transitions in dsRNA. In contrast, both antibodies robustly complexed with dsDNA and DNA–RNA hybrid substrates. Cryo-EM structural analysis confirmed that both antibodies interact with Z-NA through a conserved interaction network, and demonstrated that complexes between the antibodies and DNA-RNA hybrids adopted distinct higher-order organizations. Z22 retained binding to both Z-DNA and Z-RNA segments, whereas Z-D11 displayed substrate-dependent organization and was restricted to Z-DNA segments Molecular modelling provided mechanistic explanation for the differences in Z-RNA binding ability between the two antibodies, which was confirmed *in situ* with ADAR1-depleted, IAV-infected and JTE607-treated cells. We also demonstrate that Z-D11, in contrast to previously described Z22, is unable to induce B-to-Z-transitions in short d(CG)_6_ oligos under physiological conditions. Together, these results show that anti Z-NA monoclonal antibodies are not functionally interchangeable: whereas Z22 recognizes pre-formed Z-RNA, Z-DNA-RNA hybrids, and Z-DNA, Z-D11 recognizes Z-DNA, and DNA segments in Z-DNA-RNA hybrids. These differences are dictated by substrate composition, local geometry, and conformational accessibility, which also likely impact their ability to induce NA structural transitions. These findings have important implications for interpreting antibody-based detection of left-handed nucleic acids in biological systems.

## Introduction

Double-stranded nucleic acids can adopt alternative conformations beyond the canonical right-handed B- and A-forms, including left-handed Z-DNA and Z-RNA. These Z-conformations arise under specific sequence and environmental conditions and have been structurally characterized at high resolution ^[1,2]^, revealing distinct geometric and energetic features relative to their right-handed counterparts ^[3,4]^.

The biological relevance of Z-conformations has remained elusive for decades ^[5]^ but became clearer with the identification of proteins that selectively recognize these structures, most notably Zα domains such as those found in ADAR1 ^[6,7]^. These domains not only bind Z-nucleic acids (Z-NAs) but can also promote B-to-Z transitions, complicating their use as passive indicators of Z-conformation. Z-DNA and Z-RNA have since been implicated in a range of biological processes, particularly in immune signalling and interferon-driven responses ^[8–26]^.

This duality—recognition versus induction—poses a broader challenge for the study of Z-NAs *in vivo*, as tools used to detect Z-conformations may themselves alter the conformational landscape of nucleic acids ^[27–30]^. Monoclonal antibodies raised against Z-DNA have therefore been widely adopted as alternative probes, under the implicit assumption that they primarily bind pre-existing Z-structures without significantly perturbing nucleic acid conformation. Among these, the Z22 antibody is the most commonly used reagent for detecting Z-DNA ^[31]^, while Z-D11 has been reported to exhibit distinct binding properties, including unusually stable complexes, hysteresis, and functional effects on transcription ^[32–39]^. Despite their widespread use, it remains unclear whether these antibodies act purely as recognition elements or whether they can also contribute to Z-conformation stabilization or induction.

Recent structural studies have begun to define the molecular basis of antibody recognition of Z-DNA. High-resolution analyses of Z-D11 and Z22 in complex with Z-DNA revealed key determinants of binding, including shape complementarity to the zig-zag backbone and interactions primarily with the phosphate backbone ^[40,41]^. These findings establish a structural framework for conformational selectivity. However, these insights are derived exclusively from DNA substrates and do not address whether similar principles govern recognition of Z-RNA. Given the distinct structural and energetic properties of Z-RNA, it remains unclear whether these antibodies can engage RNA in an analogous manner, or how substrate composition influences binding specificity, selectivity, and the ability to stabilize or induce Z-conformations.

Here, we directly compare the behavior of Z22 and Z-D11 across DNA, RNA, and hybrid/chimeric substrates to dissect their specificity and capacity to engage Z-conformations. We find that both antibodies show limited interaction with CG-rich dsRNA under conditions compatible with antibody stability, indicating that they cannot induce the A-Z transition in dsRNA. In contrast, DNA–RNA hybrid substrates appear to lower the energetic barrier to Z-conformation, thereby supporting robust complex formation. Using cryo-electron microscopy, we solved the structures of Z22 and Z-D11 with the hybrid NA substrates and show that the antibodies exhibit distinct binding behaviors, including differences in substrate selectivity and complex architecture that point to fundamentally different modes of interaction. These results indicate that the two antibodies are not functionally interchangeable but instead recognize different subsets of accessible Z-like conformations defined by substrate composition, local geometry, and conformational accessibility, with important implications for interpreting antibody-based detection of left-handed nucleic acids in biological systems.

## Materials & Methods

### Cell Culture and Maintenance

Expi293 cells were maintained in OPM-293 media (OPM Biosciences) in a humidified shaking incubator at 37°C, 8% CO_2_. Primary murine embryo fibroblasts (MEFs) were generated from E14.5 C57BL/6J embryos, and immortalized using a 3T3 protocol. MEFs were maintained in DMEM supplemented with 10 % FBS (FBS, Hyclone), 1 mM sodium pyruvate, 1× GlutaMax (Thermo Fisher Scientific) and 1 % penicillin–streptomycin (Thermo Fisher Scientific), in a humidified incubator at 37 °C, 5 % CO_2_.

### Virus

IAV H1N1 strain A/Puerto Rico/8/1934 (PR8) was propagated by allantoic cavity inoculation of 10-day embryonated chicken eggs, and titres determined by standard plaque assay on Madeline Darby Canine Kidney cells (MDCK, ATCC, CCL-34).

### Protein Expression

PcDNA3.1 Z-D11 IgG heavy chain and light chain plasmids were co-transfected in equal ratio into Expi293 cells directly using homemade polyethylenimine (PEI) at 1:3 DNA:PEI ratio. 18 hours post transfection, 5% (v/v) of OPM-293 Profeed (OPM Biosciences) was added to boost expression of the target proteins. Culture media were harvested five days post transfection.

Z22 IgG heavy and light chain plasmids were prepared by replacing the Z-D11 Fab sequence with the corresponding Z22 Fab sequence (Table 1). The Z22 Fab sequence was purchased as a gene block from Integrated DNA Technologies (IDT) and cloned into the Z- D11 IgG plasmids by Infusion reaction using the primers listed in Table 2.

**Table 1.**
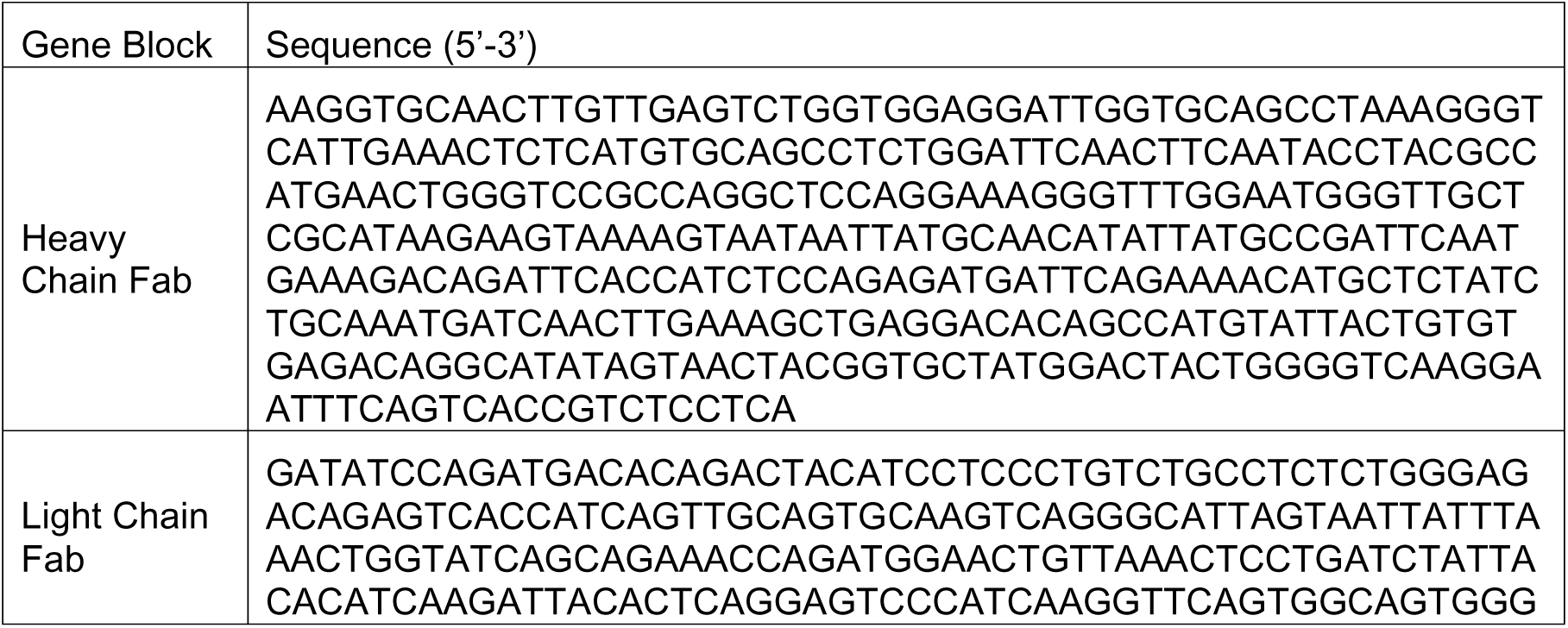
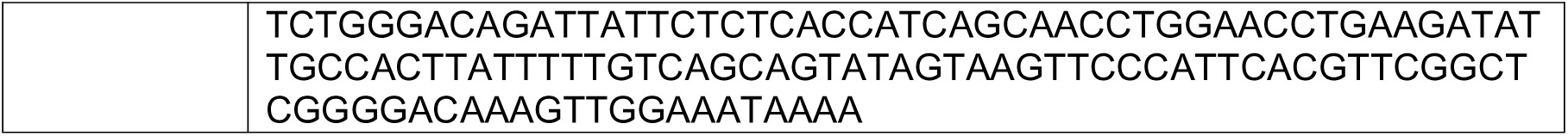
Table listing the sequence for the Z22 Heavy and Light chain Fabs used for PCR.

**Table 2.**
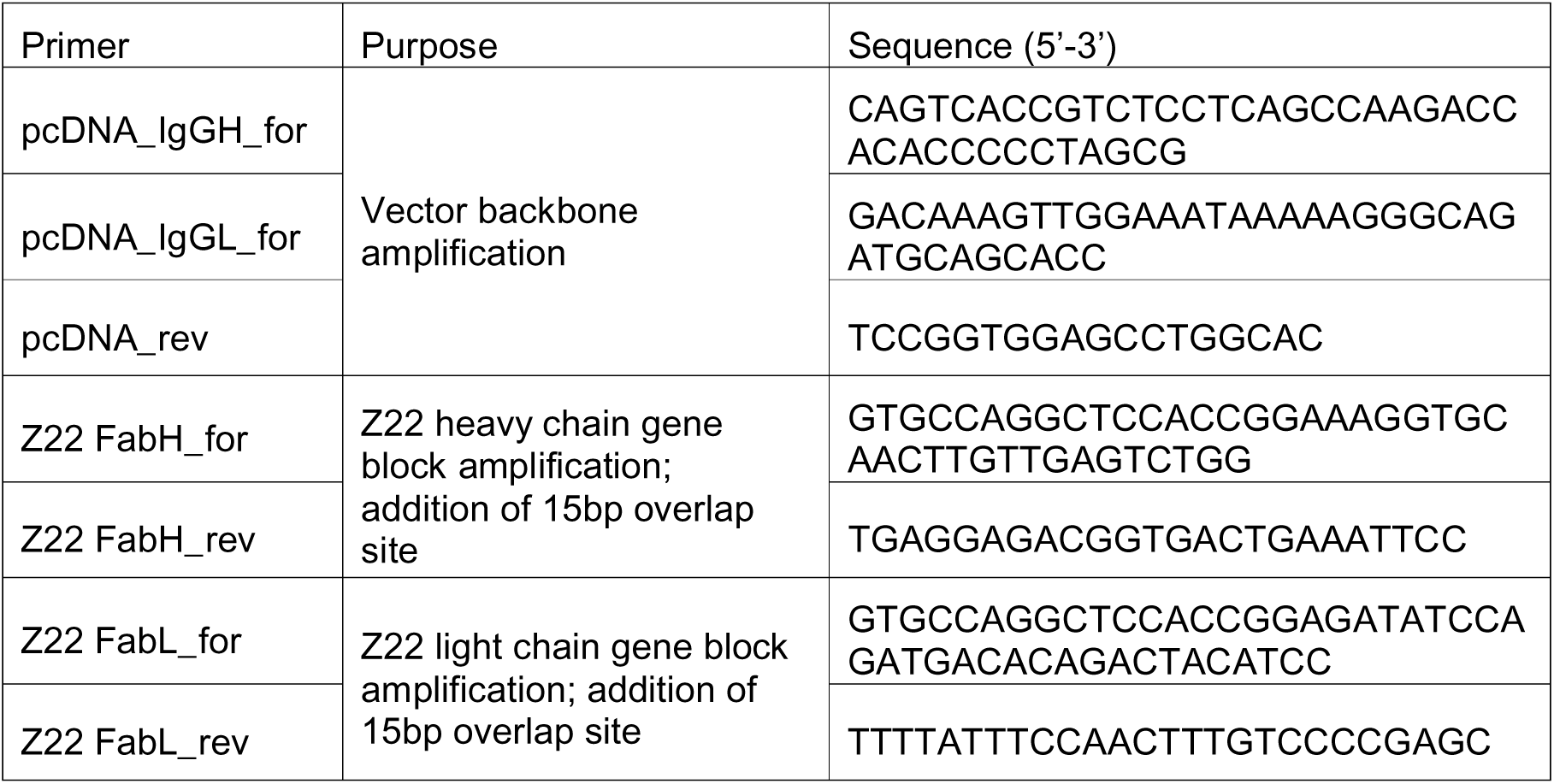
Table listing the sequence for the primers used for PCR.

### Protein Purification

In brief, harvested culture media was filtered and incubated with pre-equilibrated Protein G resin (GenScript) overnight on a roller platform at 4°C. The resin was collected in a gravity flow column and supernatant was passed through once more before washing the resin in 10 column volumes (CV) of wash buffer (1x PBS pH 7.4) twice. Antibodies were eluted from the column with 0.85 CV of elution buffer (0.1M glycine, pH 3), directly into a tube containing 15% of total volume of neutralization buffer (1M Tris, pH 8). Purified antibody was buffer exchanged into wash buffer and concentrated for downstream structural analysis.

### Cryo-EM sample preparation

The 12-mer nucleic acids (NA) were purchased from IDT. The sequence of all the (CG)_6_ NA are listed in Table 3. The hybrid-NA were resuspended in water to a final concentration of 2mM and annealed in a thermocycler using the following protocol: 95°C for 2 minutes, slow cooled to 4°C at a rate of 1°C/minute. For complexing under high salt conditions, annealed NA were then diluted to 50µM in high salt buffer (20mM HEPES pH 7.4, 2.5M NaCl, 1.25M MgCl_2_) and incubated at 37°C for 30 minutes. The antibody was added in 3-fold molar excess to the NA in high salt buffer. The resulting mixture was passed though analytical size exclusion chromatography using Superose 6 3.2/300 Increase column, pre-equilibrated in low salt buffer (20mM HEPES pH 7.4, 150mM NaCl). For complexing under low salt conditions, the antibody was mixed with the annealed NA in low salt HEPES buffer. Complexing with Tris low salt buffer was prepared according to condition reported previously ^[40]^. The mixture was subsequently passed though analytical size exclusion chromatography. Peak fractions containing the antibody and the NA were collected for cryo-grid preparation.

**Table 3.**
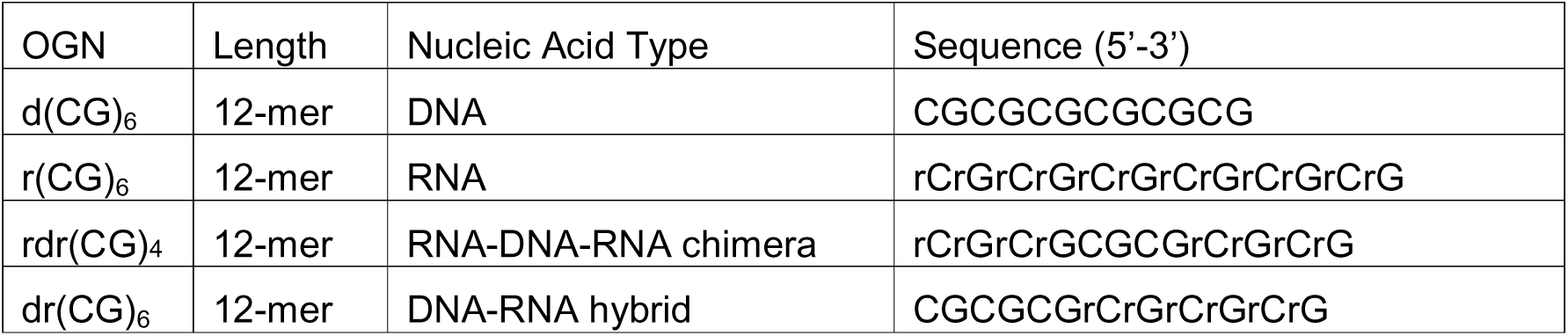
Table listing the sequence for the 12-mer NA.

### Cryo-grid preparation

Cryo-grid preparation was performed with the Vitrobot Mark IV (Thermo Fisher Scientific). Cryo-grids used were 300-mesh gold grids with R1.2/1.3 holey carbon film (Quantifoil, EMS). The grids were glow discharged for 1 minute in air plasma immediately before use. 3µL sample was pipetted onto the grid surface, blotted for 8 to 12 seconds under blot force of 2 and immediately plunge frozen in liquid ethane. The grids were subsequently clipped for data collection.

### Cryo-EM data collection and processing

All datasets of Z-D11 and Z22 complexes were collected in-house (FACTS, NTU). Micrographs were recorded on the Titan Krios electron microscope operated at 300kV. Detailed parameters for the data collection are summarized in Supplementary Table 1. The micrographs were motion corrected on CryoSPARC Live. The motion corrected micrographs were then imported into CryoSPARC-v4.7 for further processing.

For Z-D11-dr(CG)_6_ sample, particles were extracted from 5,995 micrographs. An initial model was prepared and used for Topaz training and particle extraction before 2D classification to yield 957,243 particles. A total of 294,014 particles were manually selected from the 2D classes and subjected to heterogeneous refinement using six low-pass filtered initial models (20–80 Å) to remove junk particles and resolve structural heterogeneity. One class, consisting of 270,666 particles, was re-extracted and subjected to non-uniform refinement using six low-pass filtered initial models (10–60 Å) restrained with C2 symmetry. Reference based motion correction was performed on 119,096 selected particles. These particles were further refined by non-uniform and local refinement to obtain a 3 Å map. This map was then separated into three classes by ab-initio reconstruction to obtain one good class for another round of heterogenous refinement with C1 symmetry restraints. The best class consisting of 76,537 particles, was then subjected to a final round of non-uniform and local refinement using C2 symmetry restraints to obtain a final map of the Z-D11-dr(CG)_6_ complex.

For Z22-dr(CG)_6_ sample, particles were extracted from 13,186 micrographs. An initial model was prepared and used for Topaz training and re-extraction before 2D classification. 394,629 particles were manually selected from the 2D classes for multiple iterations of ab-initio reconstruction and heterogenous refinement. 82,367 particles from the best class of heterogenous refinement was subjected to non-uniform refinement, global and local CTF refinement. Another round of non-uniform refinement followed by local refinement was performed on the same particle set to yield a 2.74 Å map showing densities of two units of a Z22 trimer surrounding the Z-NA hybrid.

For Z-D11-rdr(CG)_4_ sample, particles were extracted from 7,938 micrographs. An initial model was prepared and used for Topaz training and particle extraction, yielding 428,364 particles before 2D classification. A total of 294,014 particles were manually selected from the 2D classes and subjected to heterogeneous refinement using six low-pass filtered initial models (20–80 Å) to remove junk particles and resolve structural heterogeneity. The best class was selected for further refinement using non-uniform and global/local refinement to obtain a 2.82 Å map with 143,662 particles. This map was then separated into four classes by ab-initio reconstruction. One of the four classes was selected for another round of heterogenous, followed by homogenous, non-uniform and local refinement to obtain a final 2.88 Å map with 89,445 particles of the Z-D11-rdr(CG)_4_ complex..

For Z22-rdr(CG)_4_ sample, particles were extracted from 9,405 micrographs. An initial model was prepared and used for Topaz training and re-extraction before 2D classification. 575,960 particles were manually selected from the 2D classes for ab-initio reconstruction and heterogenous refinement. One out of six classes from the heterogenous refinement, comprising of 312,895 particles, was selected for non-uniform refinement, global and local CTF refinement. Another round of ab-initio reconstruction and heterogenous refinement was performed following the CTF refinement step to yield a good particle set of 255,330 particles. This particle set is subjected to a final round of homogeneous, non-uniform and local refinement to yield a 2.45 Å map showing densities of two units of a Z22 trimer surrounding the Z-NA chimera.

The model of the antibodies were prepared using AlphaFold 3. Each model was docked into the density map and manually refined using Coot ^[42]^ and ChimeraX ^[43]^. The model was further refined in Phenix ^[44]^ using real-space refinement with secondary structure and geometry restraints. All figures were generated by UCSF ChimeraX ^[43]^.

### Circular Dichroism

The NA was resuspended in water to a final concentration of 2 mM and annealed in a thermocycler using the following protocol: 95 °C for 2 minutes, slow cooled to 4 °C at a rate of 1 °C/minute. 4 µM of the respective NA was added into 20 mM HEPES (pH 7.4) buffer containing either salts (NaCl or MgCl_2_ or both) or other additives (Spermine or NaClO_4_) and scanned for their circular dichroism (CD) spectra. CD spectra were recorded on a Jasco-815 spectropolarimeter using 1 cm path-length quartz cuvette in a reaction volume of 500 µl at 20 °C. Scans from 220 to 320 nm were performed with 200 nm/min, 1 nm pitch and 1 nm bandwidth.

### Modelling of Z-RNA Binding to Z-D11

Z-RNA model was obtained by aligning the Fab structures of Z-D11-Z-DNA/RNA hybrid with the corresponding Z22-Z-DNA/RNA hybrid using the Matchmaker tool under UCSF ChimeraX. No further refinement against the original maps was performed to preserve the Z-DNA/RNA hybrid 3D conformation for analysis of the Z-RNA binding to Z-D11.

### Ablation of Adar gene expression by siRNA

For ablation of ADAR expression, immortalized MEFs were transiently transfected with siRNA against mouse Adar (Thermo Fisher Scientific, s211756) using Lipofectamine RNAiMAX Transfection Reagent (Thermo Fisher Scientific, 13778150). Two days after transfection with siCtrl or si*Adar*, cells were treated with IFNβ (100 ng ml⁻¹, 24 h) and examined for ADAR1 protein expression by immunoblotting or treated with IFNβ (100 ng ml⁻¹, 72 h) for immunofluorescence microscopy.

### Immunofluorescence Microscopy

CBL0137 (Cayman Chemical, Cat # 19 110), JTE-607 (MedChemExpress, Cat # HY- 110133), Anti-Z-NA Ab Clone Z22 (Absolute Antibody, Cat # Ab00783-23.0), Alexa Fluor 488 donkey anti-rabbit IgG (H + L) (Invitrogen Cat # A21206), Alexa Fluor 488 donkey anti-mouse IgG (H + L) (Invitrogen Cat # A21202), DNase I Solution (Thermo Fisher Cat # 90083) and RNase A (Thermo Fisher Cat # EN0531) were purchased from their indicated sources. Cells were plated on 8-well glass slides (EMD Millipore) and allowed to adhere for at least 24 hours before use in experiments. To detect Z-NA, cells were fixed for 10 minutes with freshly prepared 4 % (w/v) paraformaldehyde in PBS, permeabilized in 0.5 % (v/v) Triton X-100 in PBS, blocked with MAXblock blocking medium (Active Motif), and incubated overnight (16 hours) with primary antibodies at 4 °C. After three washes in PBS, slides were incubated with fluorophore-conjugated secondary antibodies for 1 hour at room temperature. Following an additional three washes in PBS, slides were mounted in ProLong Gold antifade reagent (Thermo Fisher Scientific) and imaged by Leica SP8 instrument. Fluorescence intensity was quantified using Leica LAS X software. When required, RNase A (1 mg/ml) and DNase I (25 U ml^−1^) was used for 1 hour at 37 °C before primary antibody incubation.

### Immunoprecipitation and immunoblotting

Immortalized MEFs were lysed in RIPA lysis buffer (Thermo Fisher Scientific) supplemented with protease and phosphatase inhibitors (Thermo Fisher Scientific). Cell lysates were incubated on ice for 10 min, and briefly sonicated to shear chromatin, then cleared by high-speed centrifugation (20,000g, 10 min) at 4°C. The extracts were subjected to immunoblot analysis as described previously (PMID 23613854). Sources and dilutions of primary antibodies were as follows: ADAR1 (sc-73408, Santa Cruz Biotechnology, 1:1000), GAPDH (#60004-1-Ig, Proteintech, 1:2000).

### Statistical Analysis

Statistical significance for immunofluorescence data was assessed using one-way analysis of variance (ANOVA) with either Dunnett’s multiple-comparison test. Data are presented as mean ± standard deviation (*n* = 25 cells per group). Data are representative of at least two independent experiments. *P* values of 0.05 or lower were considered to be significant.

## Results

### High thresholds for Z-formation of r(CG)_6_ dsRNA limit antibody binding

Following our analysis of Z-DNA recognition by Z-D11 and Z22, we asked whether these antibodies could also bind Z-form RNA. In principle, both antibodies might be expected to recognize Z-RNA in a manner analogous to Z-DNA. To test this, Z-D11 was incubated with a 12-mer RNA under high-salt conditions, 2.5 M NaCl supplemented with 1.25 M MgCl_2_, to promote the otherwise energetically unfavorable transition of RNA into the Z-conformation. However, analytical size-exclusion chromatography (SEC) showed no evidence of complex formation, with two distinct peaks corresponding to free RNA and free antibody (Fig. 1A). A similar SEC profile was observed when Z22 was incubated with the same 12-mer RNA (Fig. 1B), indicating that neither antibody formed a detectable complex under these conditions.

**Figure 1.**
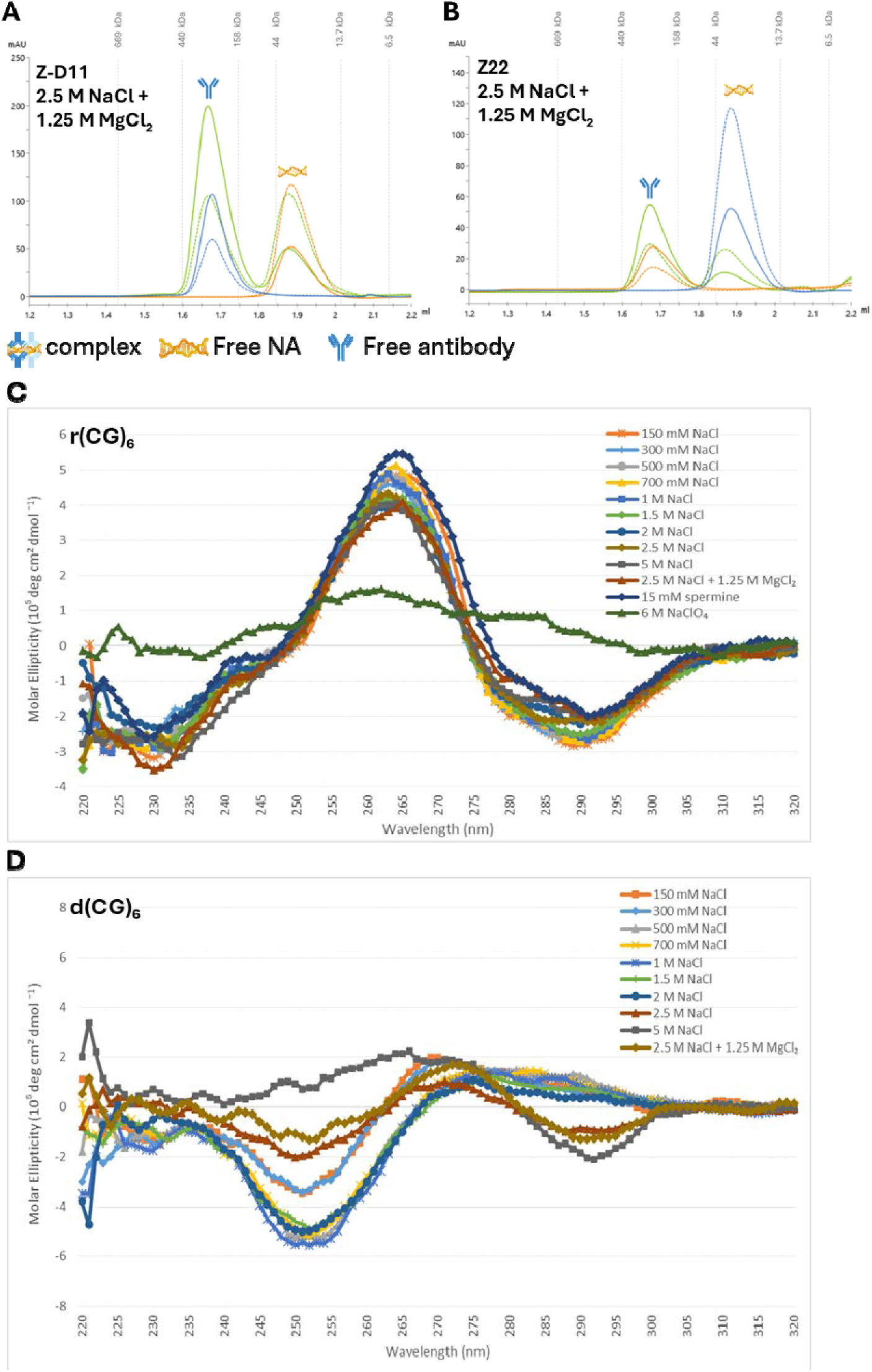
High thresholds for Z-formation of r(CG)_6_ dsRNA limits antibody binding. Size exclusion chromatograms of **A.** Z-D11 and **B.** Z22 mixed with r(CG)₆ under high salt buffer (20 mM HEPES pH 7.4, 2.5 M NaCl, 1.25 M MgCl₂). The chromatograms for the peaks are represented in green (antibody-RNA sample), blue (antibody only sample), and yellow (RNA only sample), respectively. Absorbance at 260 nm and 280 nm are presented as dashed and solid lines, respectively. Void volume for Superose 6 Increase 3.2/300 column is approximately 0.8 mL. Circular dichoism spectra of **C.** r(CG)₆ and **D.** d (CG)₆ under varying conditions of salt and additives.

We next asked whether the absence of complex formation reflected suboptimal Z-RNA induction rather than a lack of antibody recognition. To address this, we tested a range of induction conditions, including low pH, MgCl_2_ supplementation, and the polyamine spermine, either individually or in combination (Fig. S1A–S1E). Because Z-RNA has previously been observed in 6 M NaClO_4_ ^[3,45,46]^, the RNA was also pre-treated under this condition before incubation with each antibody. Z-D11 remained soluble in 6 M NaClO_4_, whereas Z22 precipitated immediately upon addition, preventing further analysis. Nevertheless, no Z-D11– RNA complex was detected following NaClO_4_ treatment (Fig. S1F). By contrast, both Z-D11 and Z22 readily formed complexes with Z-DNA under equivalent assay conditions, confirming that the SEC system was capable of detecting antibody–nucleic acid binding ^[41]^.

To determine whether the RNA had adopted the Z-conformation under the tested conditions, we performed circular dichroism (CD) spectroscopy across a range of solution conditions, including increasing NaCl concentrations from 150 mM to 5 M, 2.5 M NaCl supplemented with 1.25 M MgCl_2_, 15 mM spermine, and 6 M NaClO_4_. The CD spectra indicated that the 12-mer RNA displayed features consistent with Z-form RNA only in 6 M NaClO_4_, as shown by reduced ellipticity at approximately 265 nm and an increased signal at approximately 295 nm (Fig. 1C). In contrast, the 12-mer DNA exhibited a CD profile consistent with left-handed Z-form DNA under substantially milder conditions (Fig. 1D), consistent with the higher energetic barrier associated with conversion of RNA to the left-handed conformation ^[3]^.

Together, these results suggest that the failure to detect antibody–RNA complex formation most likely reflects the difficulty of stabilizing Z-form RNA under antibody-compatible conditions, rather than an intrinsic inability of Z-D11 or Z22 to recognize Z-RNA.

### DNA-RNA hybrid dr(CG)**_₆_** reveals differential binding modes of Z-D11 and Z22

To overcome the limitations associated with formation and stabilization of unmodified Z-form RNA, hybrid nucleic acids were employed for downstream complex formation studies. The hybrid dr(CG)_6_, comprises a 6-mer DNA linked to a 6-mer RNA. Complex assembly was performed under high-salt conditions (2.5 M NaCl and 1.25 M MgCl_2_), which promote the B-to-Z transition.

Analytical SEC confirmed the formation of higher-order oligomeric complexes with both Z- D11 and Z22, as shown by shifts in elution profiles toward higher molecular weights relative to the free components (Fig. 2A, 2D). These results indicate that the hybrid substrate adopts a Z-conformation recognizeable by both mAbs.

**Figure 2.**
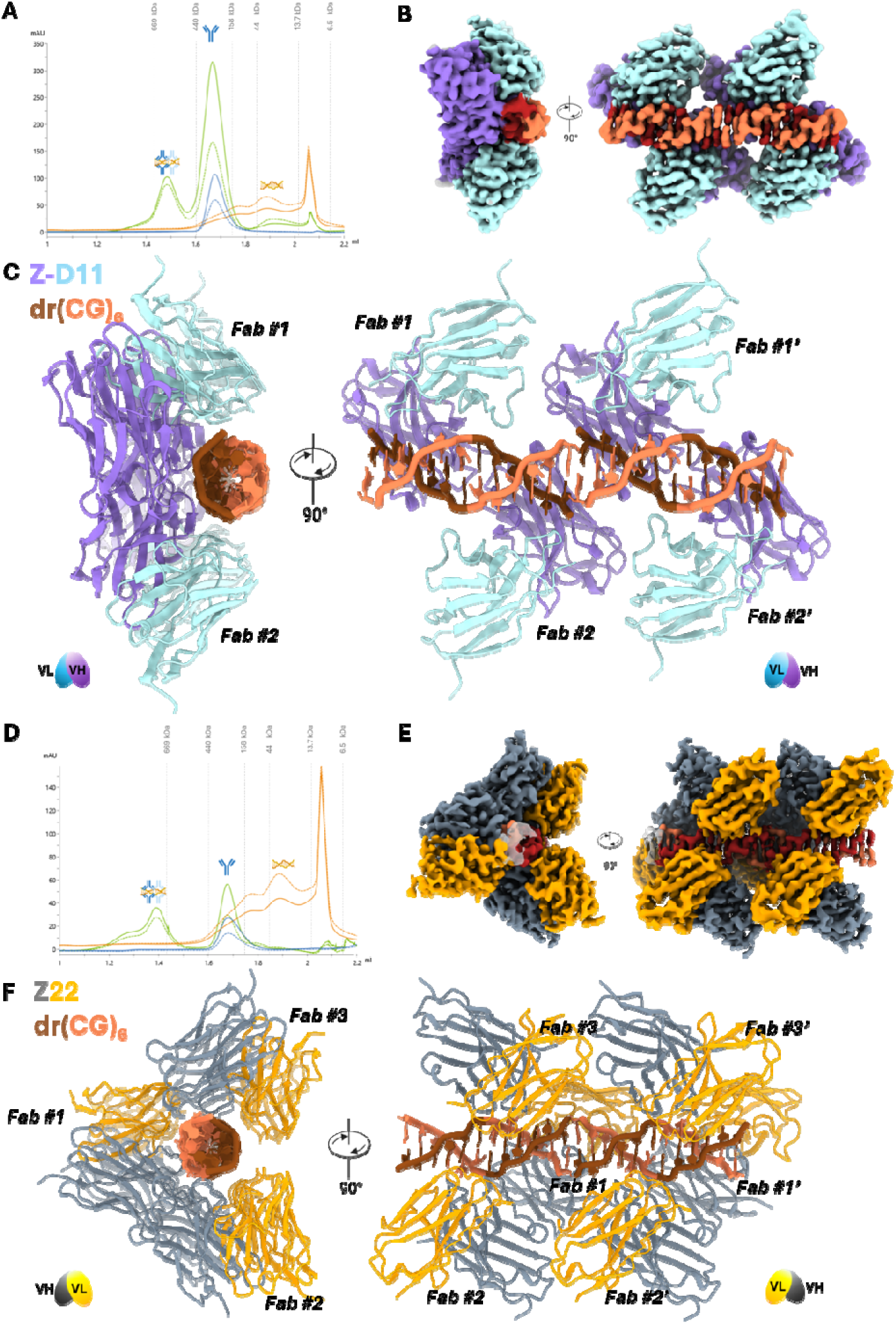
Cryo-EM structure of Z-DNA binding monoclonal antibody, Z-D11 and Z22, to DNA-RNA hybrid nucleic acids. **A.** Size exclusion chromatograms of Z-D11 with dr(CG)₆ under high salt buffer (20 mM HEPES pH 7.4, 2.5 M NaCl, 1.25 M MgCl₂). Cryo-EM density **B.** map and **C.** model of Z-D11 mAb binding to two units of dr(CG)₆. Each fab within the Z- D11 mAb-dr(CG)₆ dimer is labelled from 1 to 2, and the second trimer unit labelled from 1’ to 2’. **D.** Size exclusion chromatograms of Z22 with dr(CG)₆ under high salt buffer (20 mM HEPES pH 7.4, 2.5 M NaCl, 1.25 M MgCl₂). Cryo-EM density **E.** map and **F.** model of Z22 mAb binding to two units of dr(CG)₆. Each fab within the Z22 mAb-dr(CG)₆ trimer is labelled from 1 to 3, and the second trimer unit labelled from 1’ to 3’. Segments of the Z-DNA is coloured in brown, and Z-RNA is coloured in coral, respectively. Z-D11 is coloured in purple and turquoise for the heavy chain and light chain, respectively. Z22 is coloured in silver and orange for the heavy chain and light chain, respectively.

Comparison of SEC elution volumes further revealed differences between the antibody– hybrid nucleic acid complexes. The Z-D11–dr(CG)_6_ complex eluted at a later volume, approximately 1.48 mL, corresponding to an estimated molecular weight of 440–669 kDa. In contrast, the Z22–dr(CG) _6_ complex eluted earlier, at approximately 1.39 mL, corresponding to an estimated molecular weight greater than 669 kDa. These differences suggest that the Z-D11–dr(CG)_6_ and Z22–dr(CG)_6_ complexes differ in assembly state and/or stoichiometry.

### High-Resolution Cryo-EM Structures and Stoichiometry

The final high-resolution structures obtained by single-particle cryo-EM revealed distinct higher-order assemblies for the two antibody–hybrid nucleic acid complexes. Initial processing of the Z-D11–dr(CG)_6_ and Z22–dr(CG)_6_ datasets yielded reconstructions at 3.0 Å and 2.74 Å resolution, respectively (Fig. S2, S3). Although full-length antibodies were used for complex formation, only the Fab regions were resolved in all structures, with no interpretable density observed for the Fc regions.

The Z-D11–dr(CG)_6_ complex adopted a distinct structural arrangement. In this assembly, Z- D11 Fabs preferentially aligned along the DNA face of the hybrid duplex (Fig. 2B, 2C), producing a face-biased architecture rather than the axial organization previously observed for Z-D11 bound to Z-DNA ^[41]^ (Fig. S6A). The overall alignment displayed a root mean square deviation (RMSD) range between 0.40 Å and 0.61 Å for the Fabs, and an RMSD of approximately 0.72 Å for the nucleic acid chains when compared to Z-D11 bound to d(CG)_6_ Z-DNA (PDB 9TGN ^[41]^). Despite this asymmetry, adjacent Fabs remained sufficiently close to form inter-Fab contacts involving heavy-chain residues Gly26 and Phe27, stabilized by hydrogen bonds at approximately 3.4 Å.

In contrast, the Z22–dr(CG)_6_ complex adopted the canonical dimer-of-trimers architecture, with Fabs arranged along the duplex axis (Fig. 2E, 2F). This organization is consistent with the binding mode previously reported for Z22 in complex with Z-DNA ^[40,41]^, with the RMSD reported within the ranges of 0.25 Å to 0.47 Å across the whole structure, indicating high structural similarity between this complex and that of the Z-DNA complex (Fig. S6B). Within this assembly, adjacent Fabs formed well-defined interfaces involving residues Gly26 and Asn28, stabilized by hydrogen bonds at distances of approximately 3.7–3.9 Å. These contacts are consistent with those observed in the Z22–Z-DNA complex and support a conserved mechanism of higher-order assembly when antibody-binding sites are closely spaced.

Together, these findings support a model in which the Z-form duplex defines the overall assembly framework, while local binding geometry determines whether stabilizing Fab–Fab interfaces can form. Differences in this balance likely underlie the distinct substrate selectivity and assembly behaviour observed between Z-D11 and Z22.

### Molecular Basis for Hybrid Z-NA dr(CG)_6_ Recognition

To define the molecular basis underlying the conserved recognition and variable assembly behaviour described above, we examined the Fab–nucleic acid interfaces at residue-level resolution. Local map quality at the Fab–NA interfaces was well resolved across all complexes, enabling confident placement of side chains and identification of hydrogen-bonding and aromatic interactions (Fig. 3).

**Figure 3.**
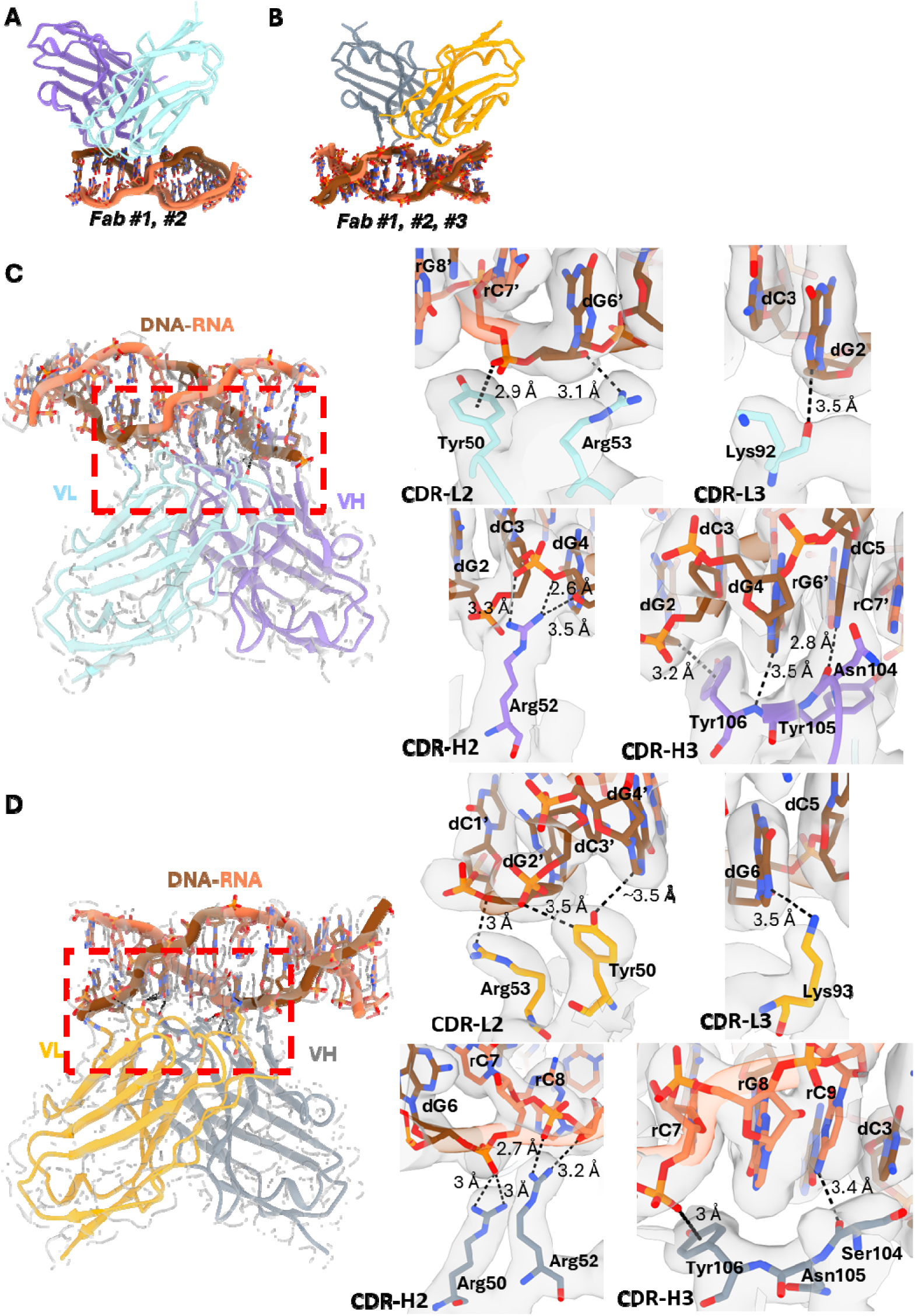
Model fitting and molecular contacts of the Z-D11 and Z22-dr(CG)_₆_-DNA complex. Alignment of the **A.** Z-D11-dr(CG)₆ and **B.** Z22-dr(CG)₆ fabs along the dr(CG)₆ axis. Local side chain fitting of **C.** Z-D11 and **D.** Z22 within the cryo-EM density map at the antibody-DNA interaction interface. Amino acid residues and the corresponding interacting nucleic acids are labelled. Segments of the Z-DNA is coloured in brown, and Z-RNA is coloured in coral, respectively. Z-D11 is coloured in purple and turquoise for the heavy chain and light chain, respectively. Z22 is coloured in silver and orange for the heavy chain and light chain, respectively.

Across both structures, Z-D11 and Z22 recognize Z-form nucleic acid through a conserved interaction network dominated by electrostatic contacts with the phosphate backbone and aromatic interactions with exposed bases. Backbone stabilization is mediated primarily by residues in CDR-H2 and CDR-L2, whereas CDR-L1, CDR-L3, and CDR-H3 contribute to base and sugar recognition through direct hydrogen bonding and π-stacking interactions.

In the Z-D11–dr(CG)_6_ complex, alignment of individual Fab subunits revealed minimal local conformational variation, with root mean square deviation (RMSD) values below 0.6 Å across 119 atom pairs (Fig. 3A). Stabilization of the phosphate backbone involves charged and polar residues, notably Arg52 in CDR-H2, as well as Tyr50 and Arg53 in CDR-L2, which form contacts ranging from 2.9 Å to 3.5 Å. Z-D11 recognition is further supported by residues in CDR-L3 and CDR-H3 positioned near exposed bases and sugar moieties in the major groove, forming contacts at distances ranging from 2.8 Å to 3.5 Å (Fig. 3D, S8A).

Similarly, Z22 engages the hybrid nucleic acid through the same overall recognition strategy. Alignment of individual Z22 Fab subunits also revealed minimal conformational differences, with RMSD values below 0.35 Å across 122 atom pairs (Fig. 3B). Backbone contacts involve key residues including Arg50 and Arg52 in CDR-H2, and Tyr50 and Arg53 in CDR-L2, with bond lengths ranging from 2.7 Å to 3.5 Å. Base and sugar interactions are mediated by Lys93 from the light chain, together with Ser104 and Tyr106 from the heavy chain (Fig. 3E, S8B).

Despite this conserved recognition interface, both hybrid substrate complexes exhibit reduced binding footprints relative to their corresponding antibody–Z-DNA complexes. In the Z-D11–dr(CG)_6_ complex, the binding footprint is reduced to approximately six nucleotides and is confined to a single face of the duplex. A comparable reduction in footprint is also observed for the Z22–dr(CG) complex, despite retention of the canonical axial binding mode.

Importantly, these substrate-dependent differences in footprint and binding continuity occur without detectable changes in Fab conformation. Alignment of the Fab regions across structures revealed RMSD values below approximately 0.6 Å across 120 atom pairs, indicating that substrate composition modulates the extent and spatial distribution of antibody binding rather than altering the underlying recognition chemistry.

Together, these results demonstrate that Z-form recognition is mediated by a conserved interface, while hybrid NA substrates impose a reduced and context-dependent binding footprint that shapes higher-order organization.

### Z-D11 and Z22 Display Differential Tolerance for Nucleic Acid Scaffolds

The inability to obtain stable antibody complexes with pure r(CG)_6_ RNA suggested that the energetic barrier to Z-RNA formation remained too high under antibody-compatible conditions. To further test whether incorporation of DNA could facilitate Z-form stabilization while retaining RNA-containing regions, we designed an additional chimeric DNA/RNA substrate, rdr(CG)_4_. The rdr(CG)_4_ chimera consists of a 4-mer DNA CG repeat flanked by 4- mer RNA CG repeats. This chimera was used to assess whether the substrate-dependent binding behaviour observed with dr(CG)_6_ reflected a broader recognition preference of the two antibodies.

Cryo-EM analysis of the Z-D11–rdr(CG)_4_ and Z22–rdr(CG)_4_ complexes yielded reconstructions at 2.83 Å and 2.45 Å resolution, respectively (Fig. S4, S5). In both complexes, the dimer-of-trimers architecture was retained (Fig. 4A–4D), indicating that higher-order assembly is primarily templated by the underlying Z-form duplex. In the Z22– rdr(CG)_4_ complex, a similar Fab–Fab interaction mode was observed, in which contiguous binding sites permit close packing of adjacent Fabs, resulting in interfaces involving Gly26 and Asn28 that are stabilized by hydrogen bonds. When compared to the Z22-d(CG)_6_ (PDB 9TGO ^[41]^), the overall structure of Z22–rdr(CG)_4_ complex reported RMSDs in the ranges between 0.25 Å to 0.5 Å (Fig. S6D). By contrast, in the Z-D11–rdr(CG)_4_ complex, Fab binding was restricted to the DNA segment flanked by RNA regions, increasing the spacing between adjacent Fabs and precluding direct inter-Fab contacts. Similarly, the Z-D11– rdr(CG)_4_ structure displayed an RMSD range between 0.22 Å to 0.32 Å across the chains for the Fabs When compared to the Z-D11-d(CG)_6_ (PDB 9TGN ^[41]^), but showed greater variability with the nucleic acids with an RMSD range between 0.64 Å to 1.32 Å (Fig. S6C).

**Figure 4.**
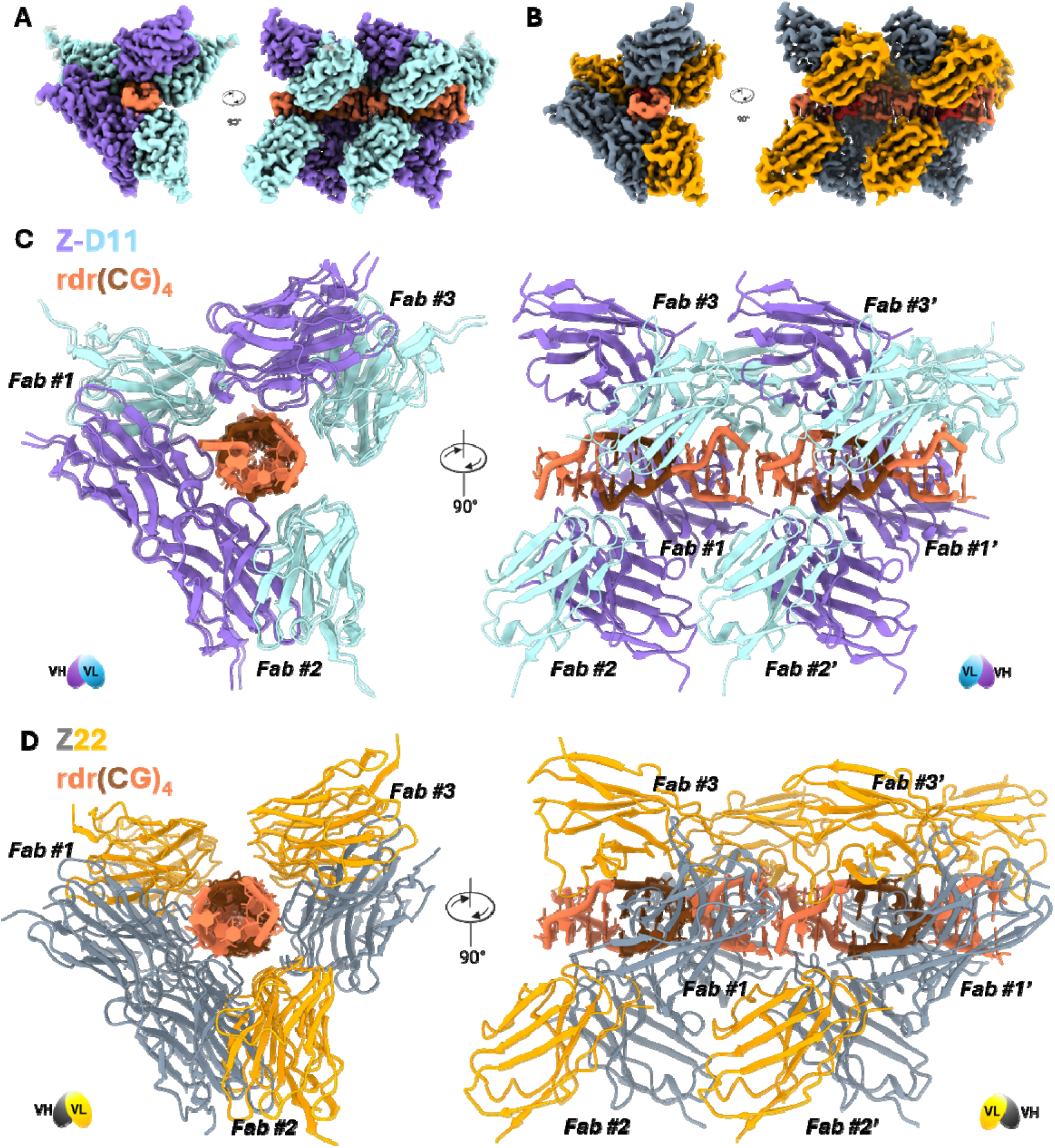
Cryo-EM structure of Z-DNA binding monoclonal antibody, Z-D11 to DNA- RNA chimera. Cryo-EM density map of **A.** Z-D11 mAb and **B.** Z22 mAb binding to two units of rdr(CG)_4_. Cryo-EM model of **C.** Z-D11 mAb-rdr(CG)_4_ and **D.** Z22 mAb-rdr(CG)_4_ complexes. Each fab within the trimer is labelled from 1 to 3, and the second trimer unit labelled from 1’ to 3’. Segments of Z-DNA is coloured in brown, and Z-RNA is coloured in coral for all complexes, respectively. Z-D11 is coloured in purple and turquoise for the heavy chain and light chain, respectively.

Alignment of individual Fab subunits revealed minimal conformational variation, with RMSD values below 0.35 Å for both complexes (Fig. S7A–S7B). Across both structures, Z-D11 and Z22 engaged Z-form nucleic acid through a conserved interaction network similar to that observed in the dr(CG)_6_ hybrid complexes, dominated by electrostatic contacts with the phosphate backbone and aromatic interactions with exposed bases. Backbone stabilization was mediated primarily by residues in CDR-H2 and CDR-L2, whereas CDR-L1, CDR-L3, and CDR-H3 contributed to base and sugar recognition through direct hydrogen bonding and π-stacking interactions, with interaction distances ranging from 2.5 Å to 3.5 Å (Fig. S7C- S7D, S8C–S8D).

Consistent with the hybrid nucleic acid structures, Z-D11 binding was restricted to the DNA-containing region of the chimera, whereas Z22 engaged the full chimeric substrate, supporting a more permissive mode of Z-form recognition. To determine the mechanism for the selective binding for Z-D11, we modelled the Z-D11 binding to Z-RNA (Fig. S9). Modelling of the Z-RNA bound to Z-D11 revealed weakened or loss of interation sites, or direct clashes with the phosphate backbone and ribose moieties. Within the light chain, contacts between Arg53 and the O4 atom of the rG6 (deoxy)ribose was weakened, with interaction distances increasing to as much as 4.6 Å. In addition, the phenolic hydroxyl group of Tyr50 was predicted to have a weakened interaction distance of 4.1 Å with the phosphate backbone of rG6 (Fig. S9C). Positioning of the strands also abolished the initial interaction between the guanosine nucleobase with Lys92, resulting in a interaction distance that was 5.4 Å with the nearest base positioned approximately 2.8 Å away (Fig. S9D). Within the heavy chain, both Arg52 and Tyr61 of CDR-H2 was predicted to clash directly with the phosphate backbone (Fig. S9E). While there was no significant difference observed with the Tyr106 peptide bond interaction with the rG10’ nucleobase, the carbonyl-O interaction distance between Asn104 and the rC11’ cytosine nucleobase was increased from 2.9 Å to 3.4 Å (Fig. S9F). These modelling results suggest that Z-D11 exhibits base- and sugar-specific constraints that bias its recognition toward DNA-containing Z-form regions. Together, the dr(CG)_6_ and rdr(CG)_4_ structures show that DNA incorporation can lower the barrier for capturing RNA-containing Z-form duplexes, while, surprisingly, revealing distinct antibody-specific recognition behaviours: Z-D11 remains biased toward DNA-containing Z-form segments, whereas Z22 can accommodate both DNA- and RNA-containing regions within a continuous Z-form scaffold.

### Immunofluorescence Demonstrates Functional Effects Consistent with Structural Model

To demonstrate that the observed Z-D11 *in vitro* binding bias toward Z-DNA-containing segments over Z-RNA is seen under physiological conditions, we performed *in situ* immunofluorescence staining on murine embryo fibroblasts (MEFs) in which Z-RNA was induced by several different approaches, employing Z22 for comparison in these studies ^[21]^. To this end, we employed three means of Z-RNA induction *in cellular*: (1) ADAR1 p150 depletion followed by Type I Interferon (IFN) stimulation as a means to induce Z-RNAs embedded in the 3’ UTRs of IFN-stimulated gene products^[21]^; (2) Influenza A virus (H1N1 strain A/PuertoRico/8/1934 (hereafter PR8) as an inducer of both viral and host-derived Z-RNAs,^[17,19]^ and (3) the CPSF3 inhibitor JTE607 as a trigger of Disruption of Transcription Termination (DoTT), a process which produces Z-RNAs by allowing aberrant transcription downstream of genes^[19]^. All three stimuli resulted in detectable Z-RNA signals in cells, when stained with the Z22 antibody, and these signals were reduced upon RNase A digestion (Fig. 5A, 5F, 5J). In contrast no detectable signal was observed for Z-D11 under the same conditions (Fig. 5C, 5H, 5L). Knockdown of ADAR1 was confirmed by immunoblotting (Fig. 5E). Notably, both antibodies stained Z-DNA, following exposure of cells to the Z-DNA inducing agent ^[21]^ (Fig. 5N, 5P). Taken together, these results demonstrate that Z22, but not Z-D11 is capable of binding Z-form RNAs under physiological conditions, whereas both antibodies detect Z-DNA.

**Figure 5.**
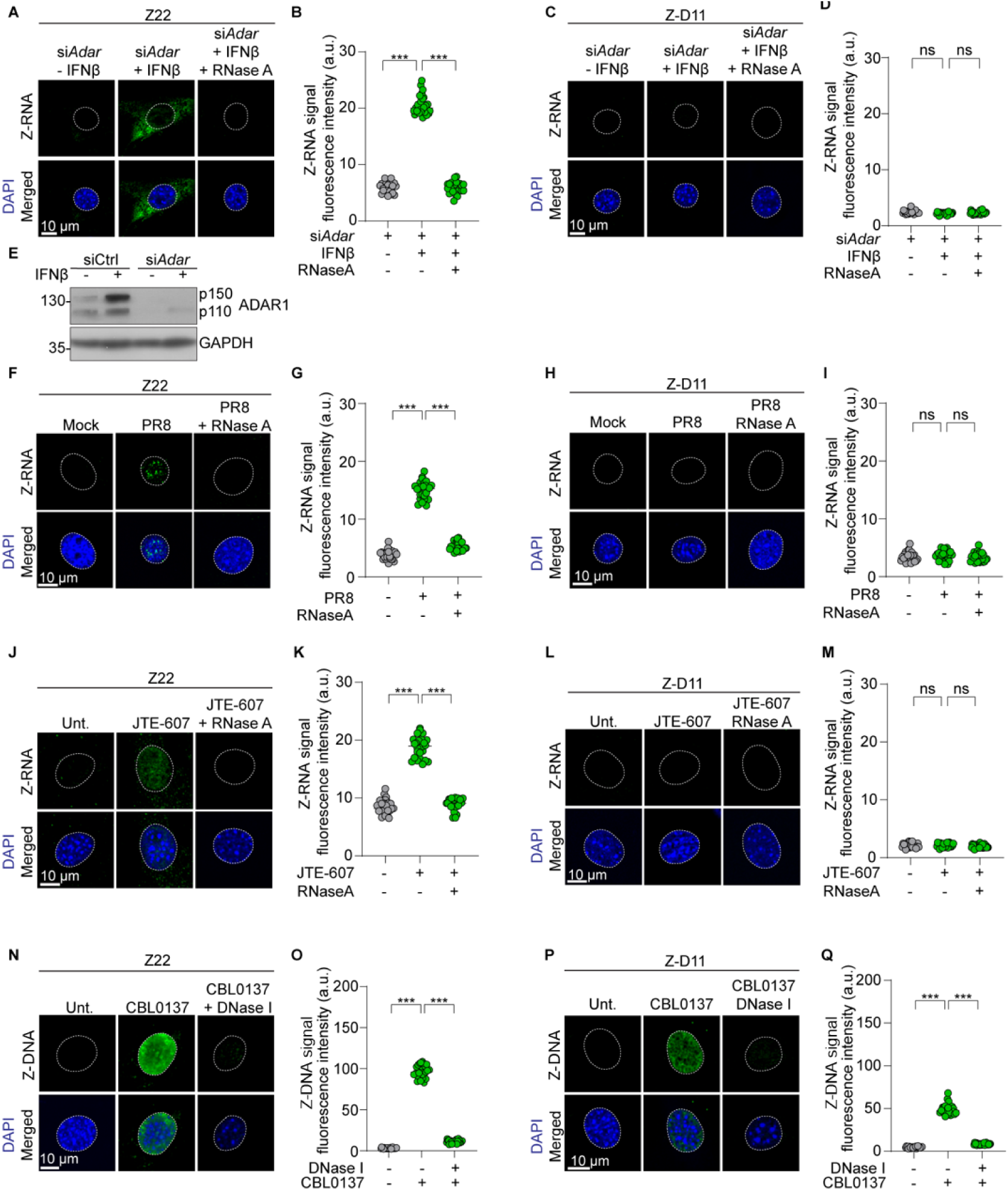
Comparison of Z-D11 and Z22 staining of Z-RNA and Z-DNA in cells. **A, C, F, H, J, L, N, P.** Immortalized MEFs were fixed after the indicated treatments and stained with rabbit monoclonal Z-NA antibodies Z22 or Z-D11 as indicated, followed by Alexa Fluor 488- conjugated secondary antibody. Nuclei were counterstained with DAPI (blue) and are outlined with dashed lines. Treatments were as follows: **A, C.** transfection of siRNAs to mouse *ADAR* (48 hrs), followed by murine IFNβ (100 ng ml⁻¹, 72 hrs); **F, H.** IAV infection (H1N1 strain PR8, MOI = 5, 8 hrs); **J, L.** murine IFNβ (100 ng ml⁻¹, 48 hrs) followed by JTE- 607 (200 μM, 48 hrs); **N, P.** CBL0137 (5 μM, 6 hrs). **E.** Levels of ADAR1 p150 and p110 isoforms in immortalized MEFs with or without IFNβ (100 ng ml⁻¹, 24 h) treatment after transfection with siCtrl or si*ADAR.* **B, D, G, I, K, M, O, Q.** Quantification of Z-RNA fluorescence intensity (arbitrary units, a.u.), corresponding to the images shown in **A, C, F, H, J, L, N, P**, respectively, are shown to the right of each set of images. Data are presented as mean ± SD (n = 25 cells per group). Statistical significance was determined by one-way ANOVA with Dunnett’s multiple-comparisons test. \*\*\**P* < 0.0005. ns, no significance. Data are representative of at least two independent experiments. Scale bar represents 10 μm.

### Z-D11 Does Not Promote Z-DNA Formation in short CG-rich DNA

We recently showed that Z-D11 does not induce B-to-Z transitions in alternating purine-pyrimidine stretches embedded in topologically relaxed plasmid DNA under physiological conditions ^[41]^. However, based on the pronounced differences between both mAb in their ability to stably bind Z-RNA segments, we decided to directly compare both antibodies in their ability to induce B-to-Z transitions in isolated short dGC oligos. To assess this, analytical SEC was performed for the antibody-DNA mixtures incubated under low-salt conditions (150 mM NaCl) in either HEPES- or Tris-based buffer. Under low-salt conditions, analytical SEC showed that Z22, but not Z-D11, forms a distinct complex with d(CG)_6_ DNA, consistent with Z-DNA induction (Fig. S10A-S10B, S10D-S10E).

We next used CD spectroscopy to evaluate the mechanism of antibody-mediated Z-DNA induction. Z-D11 or Z22 was titrated into 12-mer CG-DNA at room temperature, and CD spectra were collected at increasing antibody-to-DNA ratios. Consistent with the SEC data, addition of Z-D11 under low-salt conditions did not produce the CD spectral inversion characteristic of Z-DNA formation, even at a three-fold molar excess of antibody over DNA. Prolonged incubation of the mixture for up to 30 minutes also did not result in substantial spectral inversion (Fig. 6A). In contrast, addition of Z22 produced a modest increase in the CD signal at approximately 295 nm when added at a three-fold molar excess over DNA, indicating partial induction of Z-form DNA in solution (Fig. 6C).

**Figure 6.**
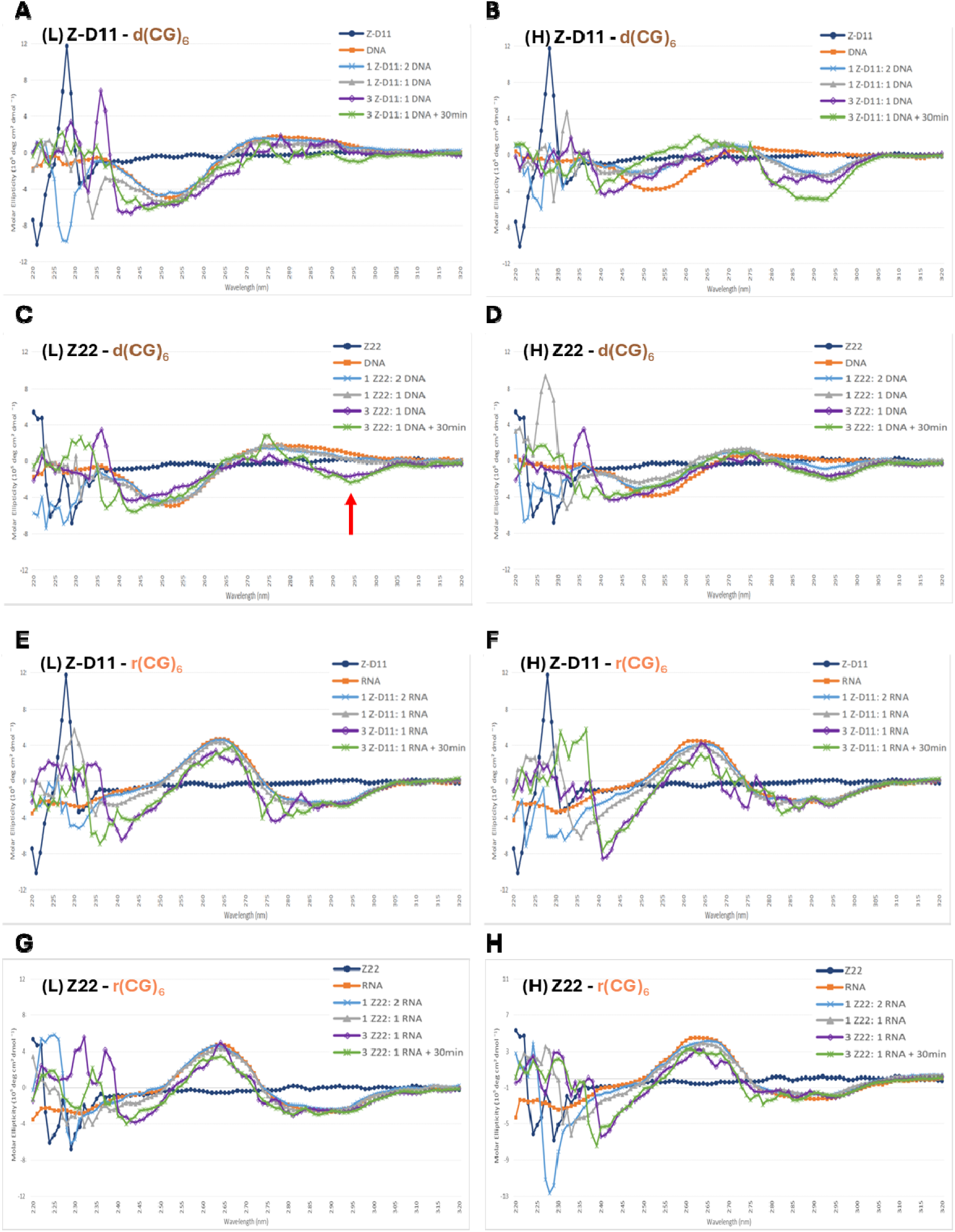
Z-D11 and Z22 have differential properties in inducing Z-formation of short CG-rich DNA. Circular dichroism spectra of d(CG)₆ in the presence of Z-D11 or Z22 under **A./C.** low salt buffer (20 mM HEPES pH 7.4, 150 mM NaCl) and **B./D.** high salt buffer (20 mM HEPES pH 7.4, 5 M NaCl); and r(CG)₆ in the presence of Z-D11 or Z22 under **E./G.** low salt buffer (20 mM HEPES pH 7.4, 150 mM NaCl) and **F./H.** high salt buffer (20 mM HEPES pH 7.4, 2 M NaCl).

Because DNA alone is unlikely to adopt the Z-conformation below 2.5 M NaCl (Fig. 1D), we next asked whether antibody binding could stabilize Z-DNA under conditions that partially favor Z-form formation. Interestingly, when CD spectroscopy was repeated in 2 M NaCl, spectral inversion was observed at lower antibody-to-DNA ratios for both Z-D11 and Z22, suggesting that both antibodies preferentially stabilize DNA in the Z-conformation once the energetic barrier to Z-DNA formation is reduced (Fig. 6B, 6D). As a positive control for Z-DNA induction, ADAR1 Zα was added to the same DNA substrate ^[27]^. At stoichiometries exceeding two molar equivalents of ADAR1 Zα per DNA, a clear inversion of the CD spectrum was observed, consistent with robust induction of the Z-conformation (Fig. S10G).

We also asked whether Z-D11 or Z22 could induce Z-form RNA. Analytical SEC and CD spectroscopy were performed using r(CG)_6_ RNA as the substrate. As expected, neither antibody formed a detectable complex with RNA under low-salt Tris buffer conditions (Fig. S10C, S10F), consistent with the high energetic barrier associated with Z-RNA formation. Likewise, addition of either antibody at up to a three-fold molar excess did not produce CD spectral inversion under either low-salt conditions or high-salt conditions, 5 M NaCl (Fig. 6E– 6H). Similarly, ADAR1 Zα, which served as a positive control for Z-DNA induction, did not induce a detectable Z-form CD signature in RNA under the same conditions (Fig. S10H). Our results could be explained with data from a recent preprint by Krall et. al., where ADAR1 Zα converts r(CG)_6_ at three-fold slower rates than d(CG)_6_ and ten-fold slower when compared to the shorter r(CG)_3_ RNA at 42 °C ^[47]^, further supporting the conclusion of a high energetic and kinetic barrier to Z-RNA formation compared to that of Z-DNA under the conditions tested, limiting protein-mediated induction of the Z-RNA conformation. Together, these findings suggest that neither Z22 nor Z-D11 can induce the A-Z transition in dsRNA within a time-frame of < 1 hour, and establish the structural framework for understanding how Z-form geometry, substrate composition, and antibody binding preferences together shape higher-order assembly.

## Discussion

In this study, we compared the recognition mode of Z-D11 and Z22 mouse monoclonal antibodies across DNA, RNA, and hybrid/chimeric substrates to determine whether they act as equivalent probes for detecting Z-conformations. Using a panel of methods, we confirmed that both antibodies, which are of the IgGgamma2b subtype, recognize Z-form nucleic acids through a broadly conserved local interaction framework ^[40, 41]^. However, this conserved recognition chemistry does not translate into equivalent substrate tolerance or higher-order assembly behaviour. Instead, Z-D11 and Z22 differ substantially in how they respond to RNA-containing substrates, with Z-D11 showing dependence on DNA-containing Z-form regions, whereas Z22 displays tolerance for RNA-containing Z-form scaffolds. This distinction was most evident in the DNA-RNA hybrid and chimera substrates, where both antibodies formed stable complexes but exhibited differential higher-order organisations. In contrast, neither antibody was capable of forming stable complexes with r(CG)_6_ RNA.

A central conclusion from our work is that the structural compatibility with the Z-conformation does not equate to experimental accessibility. While molecular modelling suggested that both antibodies could, in principle, recognize and interact with Z-RNA, the biochemical data demonstrated that r(CG)_6_ was difficult to induce in the Z-state under conditions that preserved antibody function. SEC did not detect stable antibody-RNA complexes, and CD spectroscopy similarly showed limited evidence of Z-RNA formation under conditions used for binding. Z-RNA characteristic spectral features were only observed under extreme, non-physiologically-relevant conditions. The inability of ADAR1 Zα to induce a detectable Z-RNA CD signature under comparable conditions further support the fact that r(CG) ^[3,48][3,48]^. In this respect, the hybrid and chimeric DNA-RNA substrates were especially informative, because they provided tractable scaffolds for Z-RNA stabilization, allowing antibody binding to RNA-containing Z-form contexts to distinguish between two related but distinct features: Z-conformation accessibility and productive antibody engagement.

The structures provide a basis for differential substrate tolerance of Z-D11 and Z22. Across all structures, both antibodies engage the Z-form duplex through phosphate backbone contacts and aromatic or base-proximal interactions that read out the zig-zag geometry of left-handed helices. However, incorporation of RNA changes the local surface presentation, base-pair positioning relative to the helix axis, groove architecture and continuity of antibody binding footprint ^[3, 7]^. Z22 retains a more canonical axial dimer-of-trimers arrangement observed in pure Z-DNA complexes, consistent with broader substrate tolerance. By contrast, Z-D11 displayed greater substrate-dependent variation, most clearly demonstrated in the dr(CG)_6_ complex where the Fabs align only along the DNA face of the duplex. This suggests that Z-D11 does not simply recognize left-handedness as a general geometric feature, but requires a specific presentation of the Z-form surface compatible with a DNA-associated binding footprint. These findings indicate that the two antibodies likely recognize different subsets of accessible Z-states, consistent with our immunofluorescence data, in which both antibodies exhibited distinct staining patterns upon drug-induced Z-RNA formation or viral infection.

These findings provide direct implications for the use of antibody-based reagents to study left-handed NA. First, Z22 and Z-D11 should not be treated as interchangeable probes. Second, positive signal from either antibody should not be automatically interpreted as passive capture of pre-existing Z-conformation, because as demonstrated previously ^[40]^ and in our current study, Z22 can promote Z-form formation. Under physiologically relevant conditions, Z22, but not Z-D11, formed a detectable complex with d(CG)₆ DNA and produced a modest CD signature at high antibody excess consistent with antibody-induced Z-DNA formation. Under partially Z-favouring conditions, both antibodies stabilized Z-form DNA, indicating that antibody binding can shift the conformational equilibrium of short CG-rich DNA substrates likely due to high complex stability ^[37]^. In this context, our current and previous study ^[41]^ revealed that the Z22-obtained IF signal in CBL0137-treated cells was consistently about 2-fold stronger than that observed with Z-D11, even at much lower Z22 concentrations. One possible explanation is that Z22, in addition to recognizing pre-existing Z-DNA and shifting of DNA conformational equilibrium as shown with Z-D11, is able to induce wide-spread genomic B-to-Z transitions under favorable, drug-induced DNA topological changes. Future comparisons between Z22 and Z-D11 or engineered derivatives thereof could help to elucidate mechanisms responsible for antibody-induced B-to-Z transitions.

A negative signal, particularly with RNA-rich substrates, does not necessarily indicate the absence of Z-conformation. Rather, it may reflect a combination of high energy barrier to Z-RNA formation and antibody-specific requirements for a compatible binding footprint. In that sense, positive Z-RNA signal in cells stained with Z22 should be interpreted as detection of accessible, antibody-compatible Z-RNA states rather than definitive evidence of Z22 induction *in situ*. Given the high energetic costs of the RNA A-Z transition, antibody-induced Z-RNA formation is likely more constrained than antibody-induced Z-DNA formation. Together, these results suggest that the practical value of these antibodies lies in the reporting of Z-conformations that are both structurally accessible and geometrically compatible with the antibody, rather than providing a complete view of all possible left-handed NA states. This distinction is important for interpreting Z-NA detection in biological settings such as transcription-associated stress, chromatin dynamics, and immune signalling, and provides a foundation for developing next-generation probes with improved control over substrate bias, perturbation, and discrimination between DNA- and RNA-containing Z-states.

## Acknowledgements

This research is supported by the Singapore Ministry of Education under its Singapore Ministry of Education Academic Research Fund Tier 3 (MOET32023-0003) and the Education Academic Research Funds Tier 1 (RT22/23 and RG84/21) to DL. Work in the S.B. lab is supported by NIH R01 grants AI135025, AI144400, AI161624, and CA168621. Additional support is provided by NIH Cancer Center Support Grant P30CA006927.

We thank the scientific facility support from NTU Institute of Structural Biology and Protein Product Platform. We thank Prof. Phan AT and lab members for loaning the CD spectrophotometer. We thank members of the DL lab for their support.

## Author Contributions

DHRC, PD and DL designed the study; DHRC, YL and DL performed the biophysical and structural experiments; ZH, CY, RMW, SB carried out immunofluorescence studies; All authors participated in data analysis. DHRC, YL, PD, and DL wrote the manuscript with input from all authors.

## Data And Code Availability

The SPA-cryoEM density map of the Z-D11-dr(CG)_6_ and Z-D11-rdr(CG)_4_ complex has been deposited in EM Database under the accession code EMD-81386 and 81320, respectively. The corresponding atomic coordinates have been deposited in the Protein Data Bank under accession code 27SK and 27PG, respectively. The SPA-cryoEM density map of the Z22- dr(CG)_6_ and Z22-rdr(CG)_4_ complex has been deposited in EM Database under the accession code EMD-81339 and 81340, respectively. The corresponding atomic coordinates have been deposited in the Protein Data Bank under accession code 27QC and 27QD, respectively.

