## Supplementary Information for "Anti-Z-NA antibodies can distinguish accessible Z-form states across DNA, RNA, and DNA–RNA substrates"

**Supplementary Materials**

Supplementary Figures S1-S10

Supplementary Table 1

Supplementary Movies S1-S9

**Supplementary Figure S1.** **Optimization of Z-form induction conditions of r(CG)_6_ dsRNA.** Size exclusion chromatograms of Z-D11 in complex with r(CG)_6_ under **A.** low NaCl (150 mM NaCl), **B.** high NaCl (5 M NaCl), **C.** high MgCl₂ (1.25 M MgCl₂), **D.** acidic buffer with high MgCl₂ (50 mM MES pH 3, 1.25 M MgCl₂) **E.** 150 mM NaCl with 15 mM Spermine, **F.** 6 M NaClO_4_. All buffers contain 20 mM HEPES pH 7.4 as the buffering agent, unless otherwise stated. The chromatograms for the complex, free antibody, and free NA are represented in green, blue, and yellow, respectively. Absorbance at 260 nm and 280 nm are presented as dashed and solid lines, respectively. Void volume for Superose 6 Increase 3.2/300 column is approximately 0.8 mL.

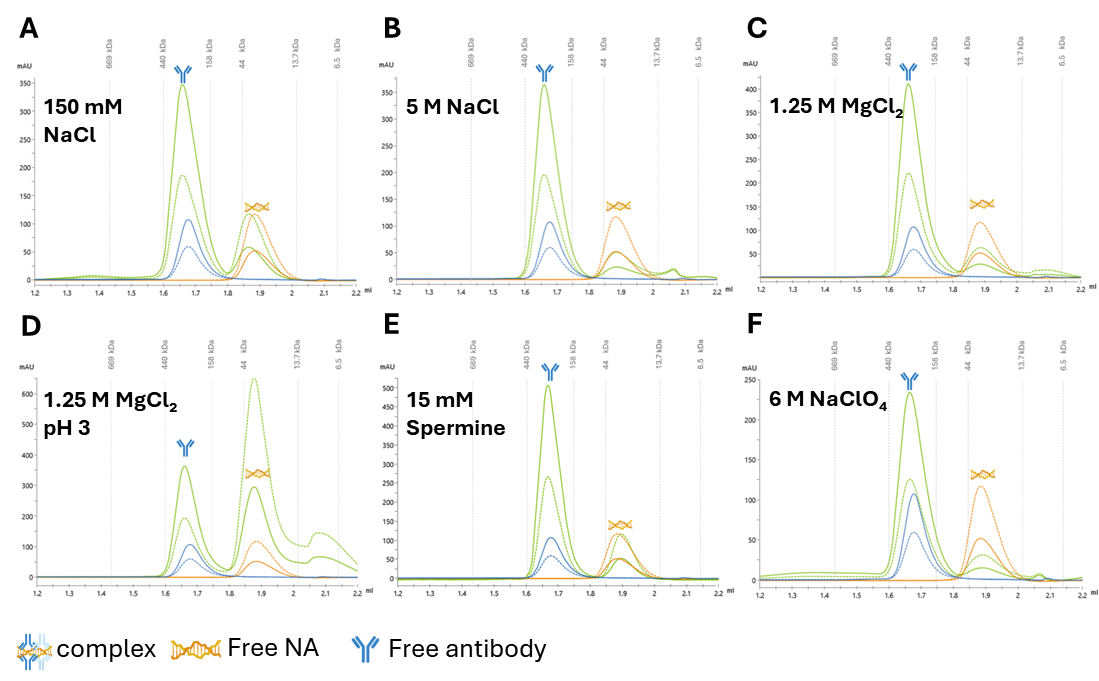

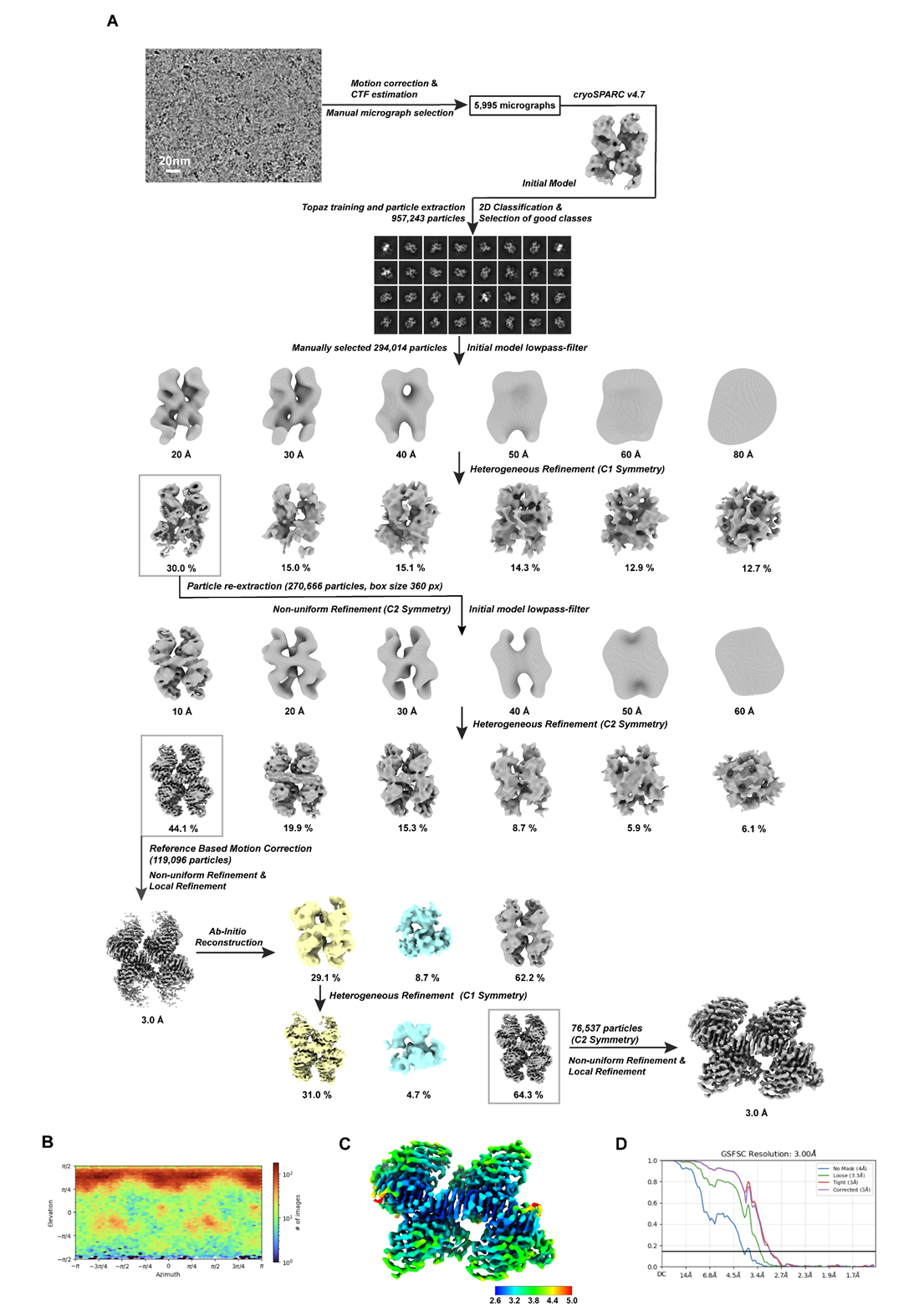
**Supplementary Figure S2.** **Structure determination of the Z-D11-dr(CG)_6_-hybrid complex.  A.** Workflow for the determination of the structure Z-D11-drd(CG)_6_. **B.** Angular distribution of particles used for the final 3D reconstruction. **C.** Local resolution distribution of the final map. **D.** FSC curves of non-uniform 3D refinement obtained from cryoSPARC.

**Supplementary Figure S3.** **Structure determination of the Z22-dr(CG)_6_-hybrid complex.  A.** Workflow for the determination of the structure Z22-drd(CG)_6_. **B.** Angular distribution of particles used for the final 3D reconstruction. **C.** Local resolution distribution of the final map. **D.** FSC curves of non-uniform 3D refinement obtained from cryoSPARC.

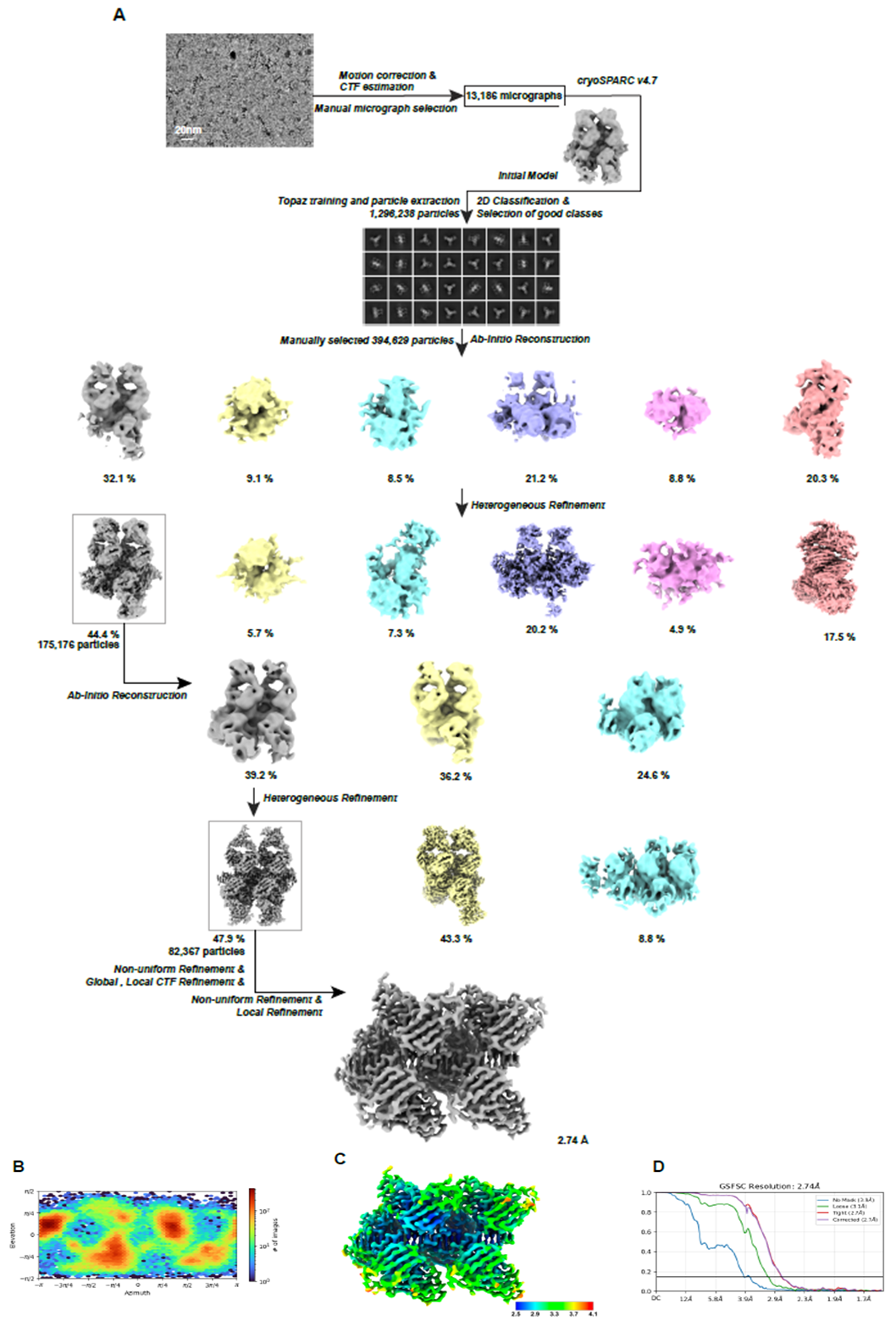

**Supplementary Figure S4.** **Structure determination of the Z-D11-rdr(CG)_4_-hybrid complex.  A.** Workflow for the determination of the structure Z-D11-rdr(CG)_4_. **B.** Angular distribution of particles used for the final 3D reconstruction. **C.** Local resolution distribution of the final map. **D.** FSC curves of non-uniform 3D refinement obtained from cryoSPARC.

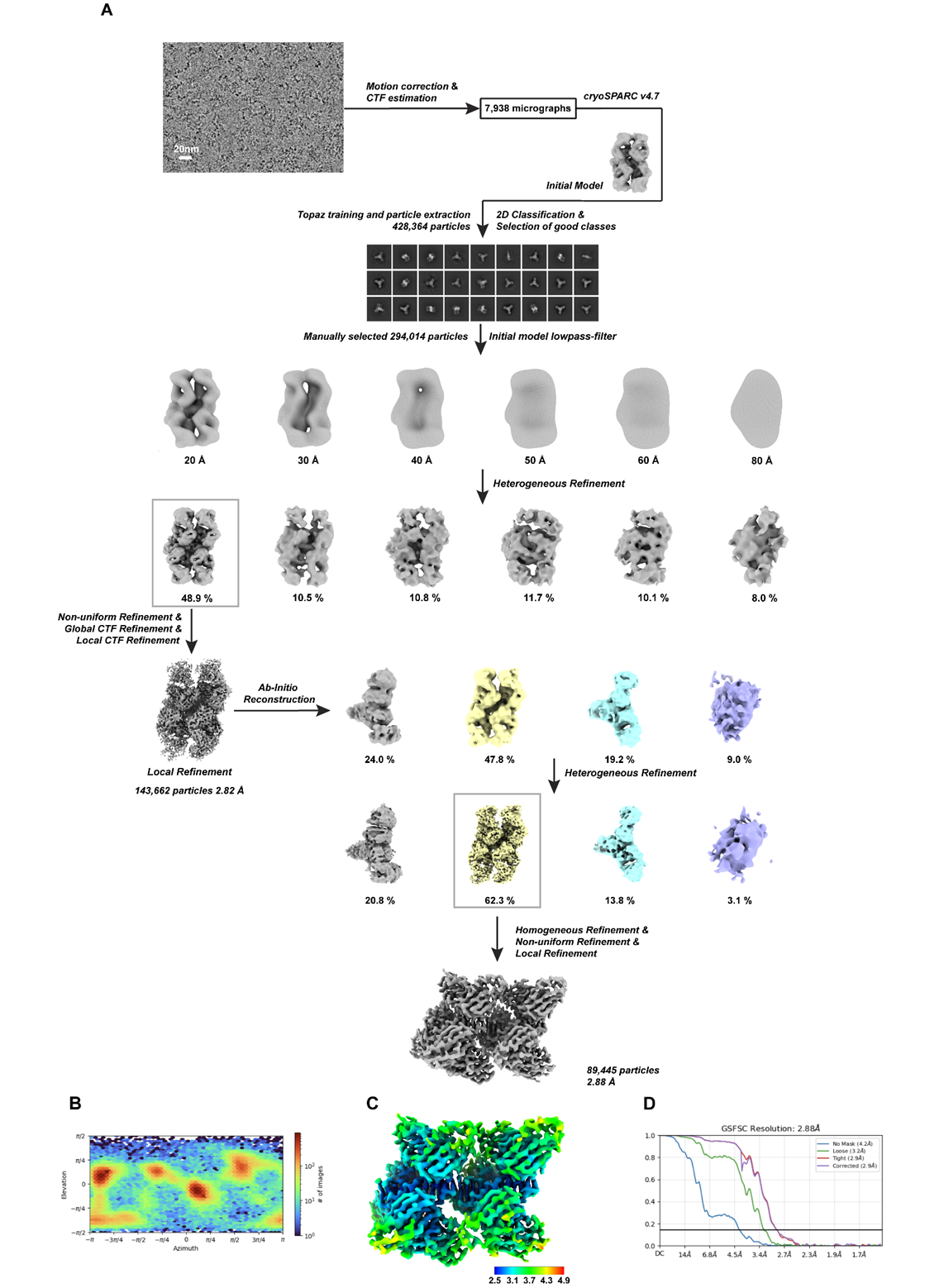

**Supplementary Figure S5.** **Structure determination of the Z22-rdr(CG)_4_-hybrid complex.  A.** Workflow for the determination of the structure Z22-rdr(CG)_4_. **B.** Angular distribution of particles used for the final 3D reconstruction. **C.** Local resolution distribution of the final map. **D.** FSC curves of non-uniform 3D refinement obtained from cryoSPARC.

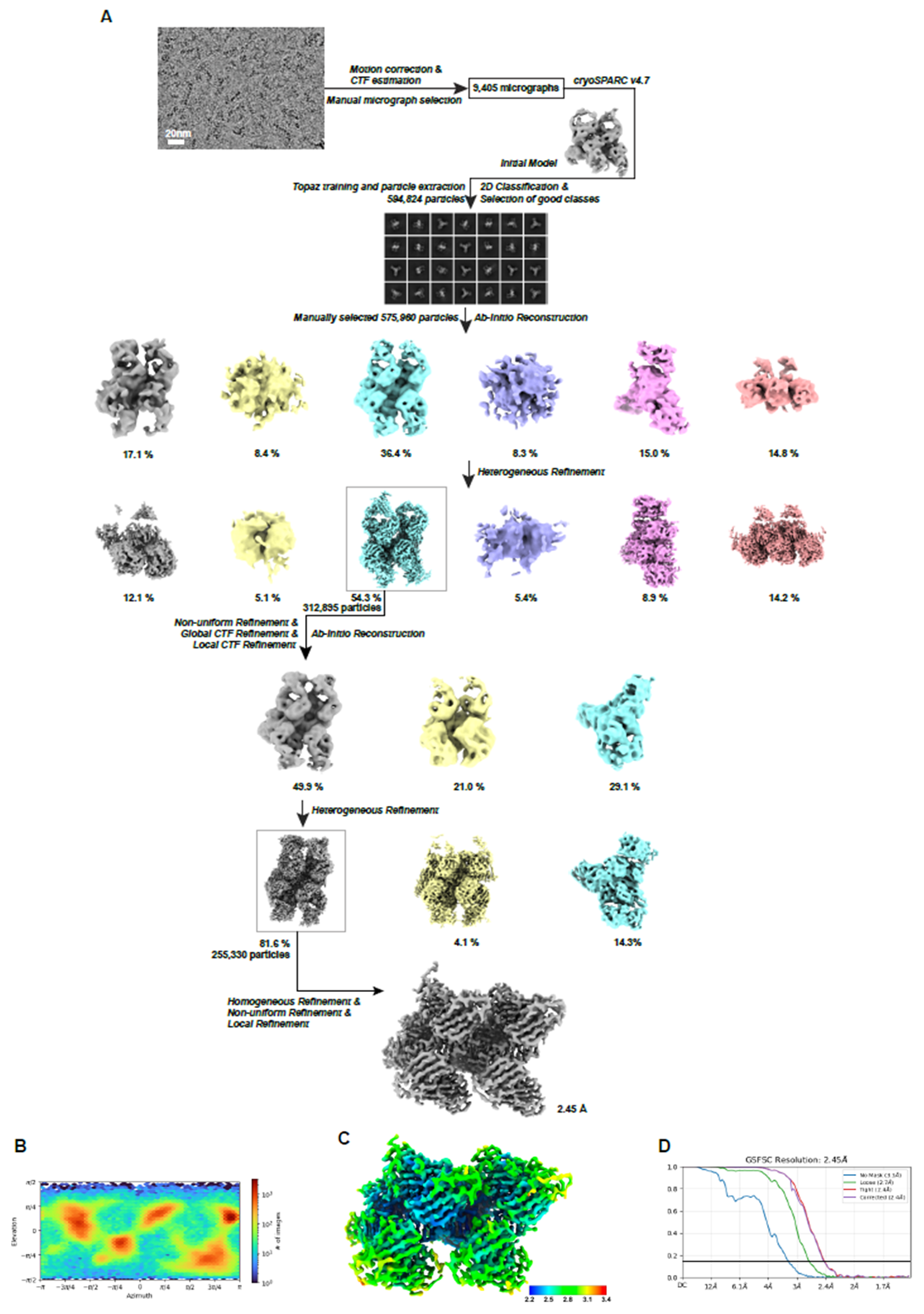

**Supplementary Figure S6. Model superposition of the dr(CG)_6_ hybrid, and rdr(CG)_4_ chimera against d(CG)_6_ Z-DNA complex.** Alignment of the **A.** Z-D11-dr(CG)_6_ and **C.** Z-D11-rdr(CG)_4_ models against the Z-D11-d(CG)_6_ complex (PDB 9TGN). Alignment of the **B.** Z22-dr(CG)_6_ and **D.** Z22-rdr(CG)_4_ models against the Z22-d(CG)_6_ complex (PDB 9TGO). DNA is coloured in brown (hybrid and chimera complexes) and lime green (Z-DNA complex); RNA is coloured in coral. Z-D11 is coloured in purple and turquoise for the heavy chain and light chain, respectively. Z22 is coloured in silver and orange for the heavy and light chain, respectively.

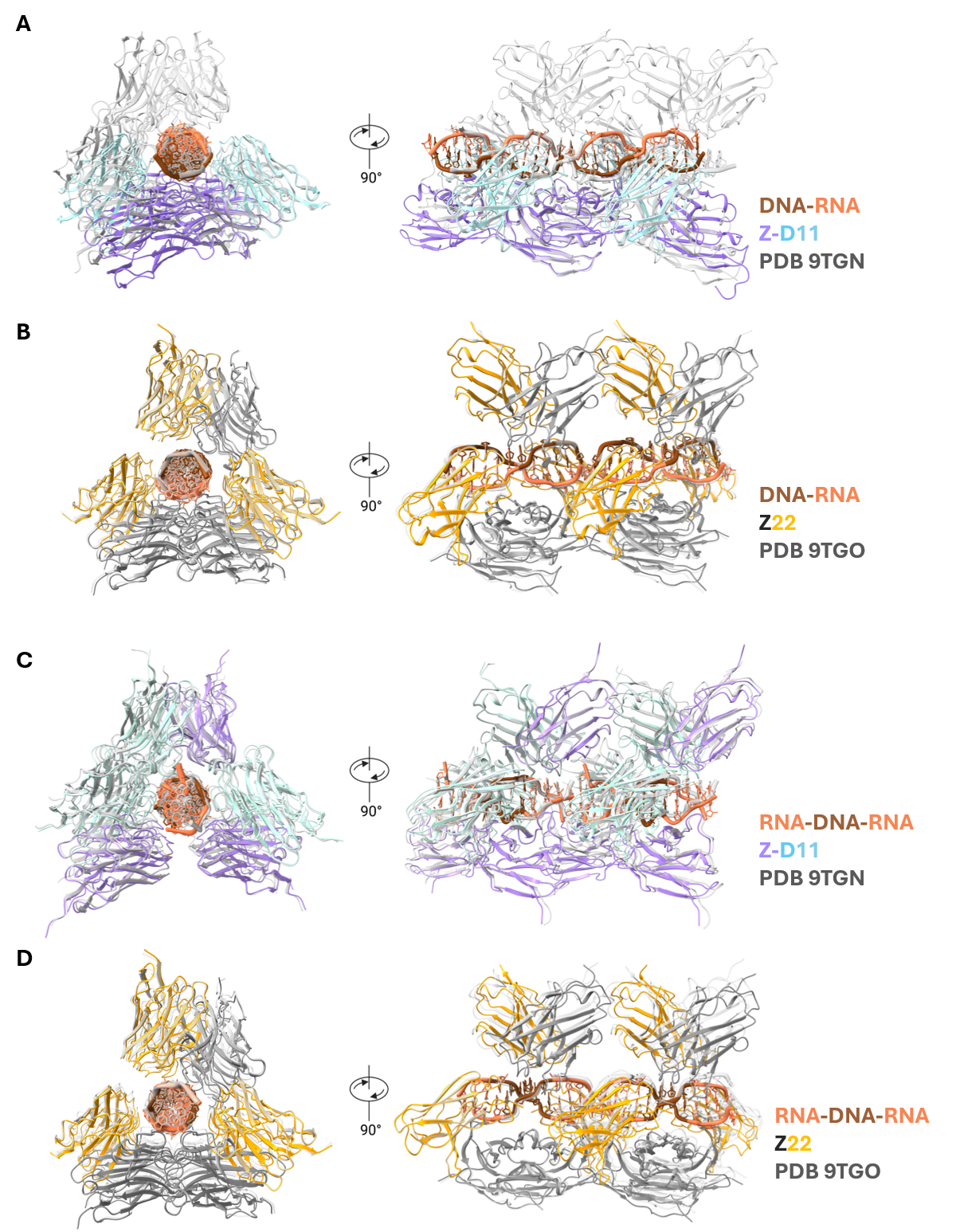

**Supplementary Figure S7. Model fitting and molecular contacts of the Z-D11-rdr(CG)_4_ and Z22-rdr(CG)_4_ chimera complex.**  Alignment of the **A.** Z-D11-rdr(CG)_4_ and **B.** Z22-rdr(CG)_4_ fabs along the rdr(CG)_4_ axis. Local side chain fitting of **C.** Z-D11 and **D.** Z22 within the cryo-EM density map at the antibody-DNA interaction interface. Amino acid residues and the corresponding interacting nucleic acids are labelled. Segments of Z-DNA is coloured in brown, and Z-RNA is coloured in coral for all complexes, respectively. Z-D11 is coloured in purple and turquoise for the heavy chain and light chain, respectively. Z22 is coloured in silver and orange for the heavy chain and light chain, respectively.

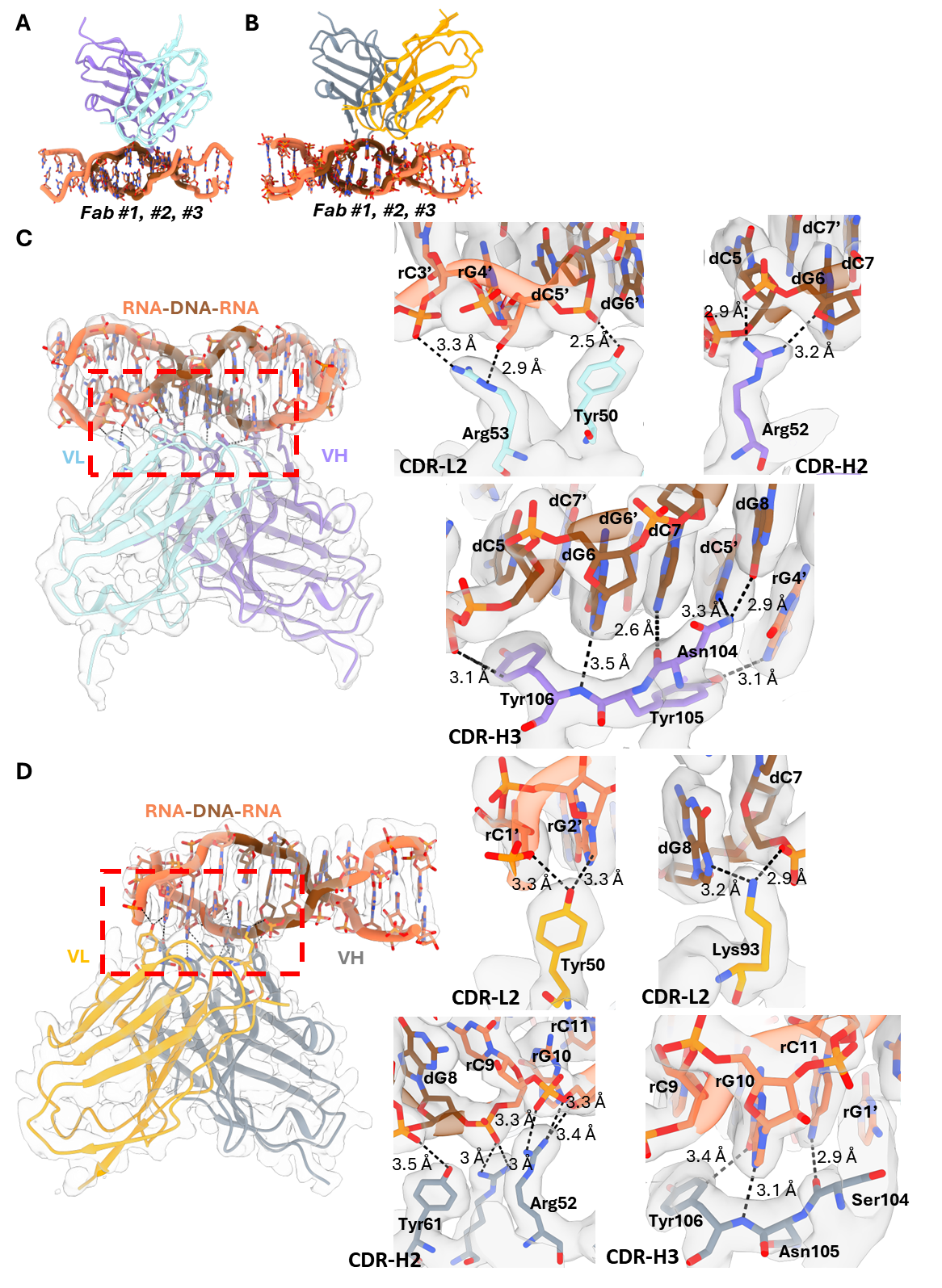

**Supplementary Figure S8. Z-D11 and Z22 utilize a conserved framework to bind Z-NA.** Cartoon representations of the interaction between **A.** Z-D11 and dr(CG)_6_, **B.** Z22 and dr(CG)_6_, **C.** Z-D11 and rdr(CG)_4_, and **D.** Z22 and rdr(CG)_4_.

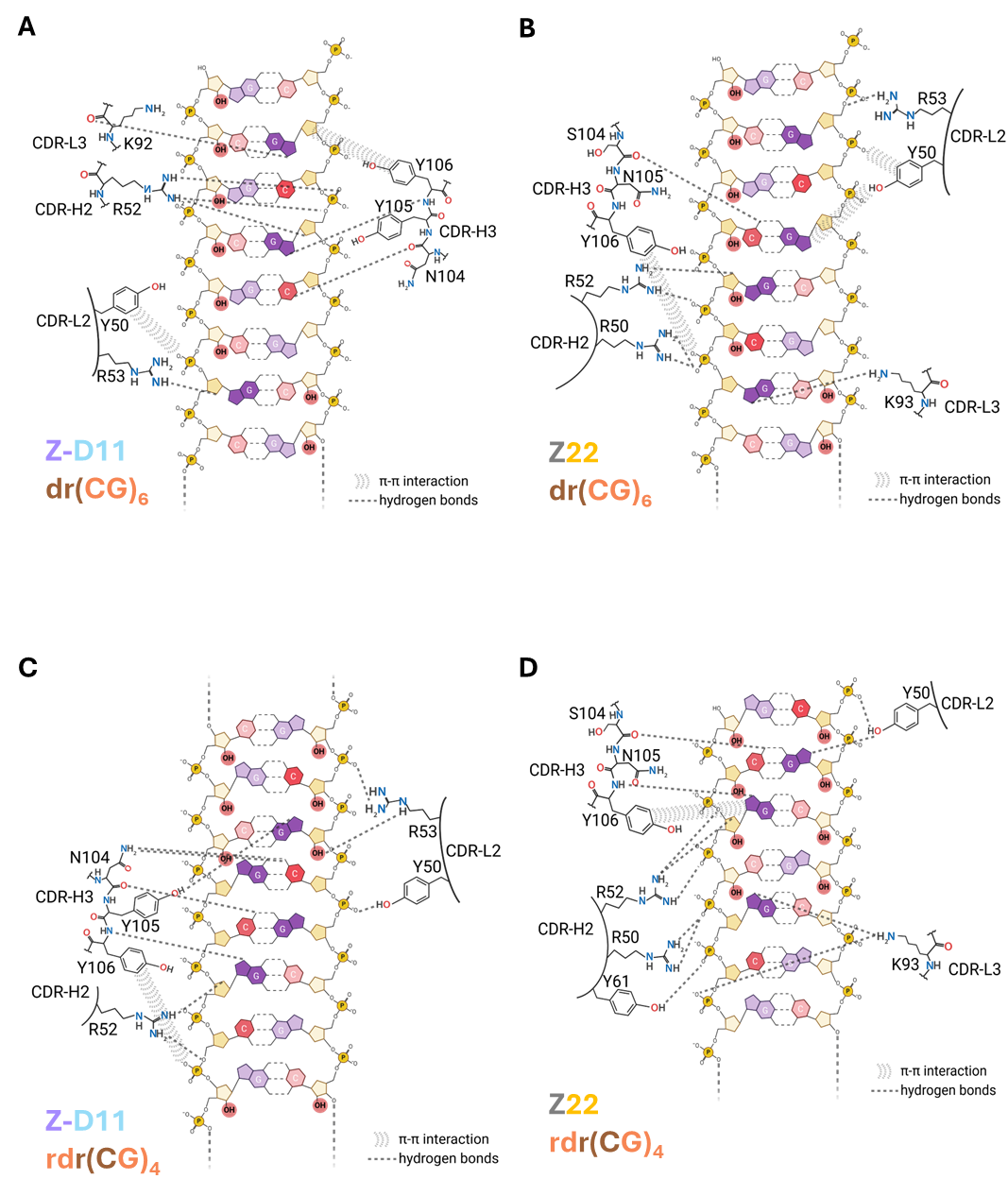

**Supplementary Figure S9. Modelling of Z-D11 binding to Z-RNA. A.** Overlay of the model of Z-D11 and dr(CG)_6_ (brown-coral) against the model of Z22 and dr(CG)_6_ (lime green-teal). **B.** Alignment of an individual Fab along the dr(CG)_6_ axis. Interaction interface between Z-D11 and the NA within **C.** CDR-L2, **D.** CDR-L3, **E.** CDR-H2, and **F.** CDR-H3. Black dotted lines represent the original interaction between Z-D11 and Z-DNA; Yellow lines represent the predicted interaction between Z-D11 and Z-RNA. Atoms in direct clash with each other are depicted as balls and boxed in red. DNA is coloured in brown (original orientation) and lime green (rotated orientation); RNA is coloured in coral (original orientation) and teal (rotated orientation). Z-D11 is coloured in purple and turquoise for the heavy chain and light chain, respectively. Z22 is coloured in gray.

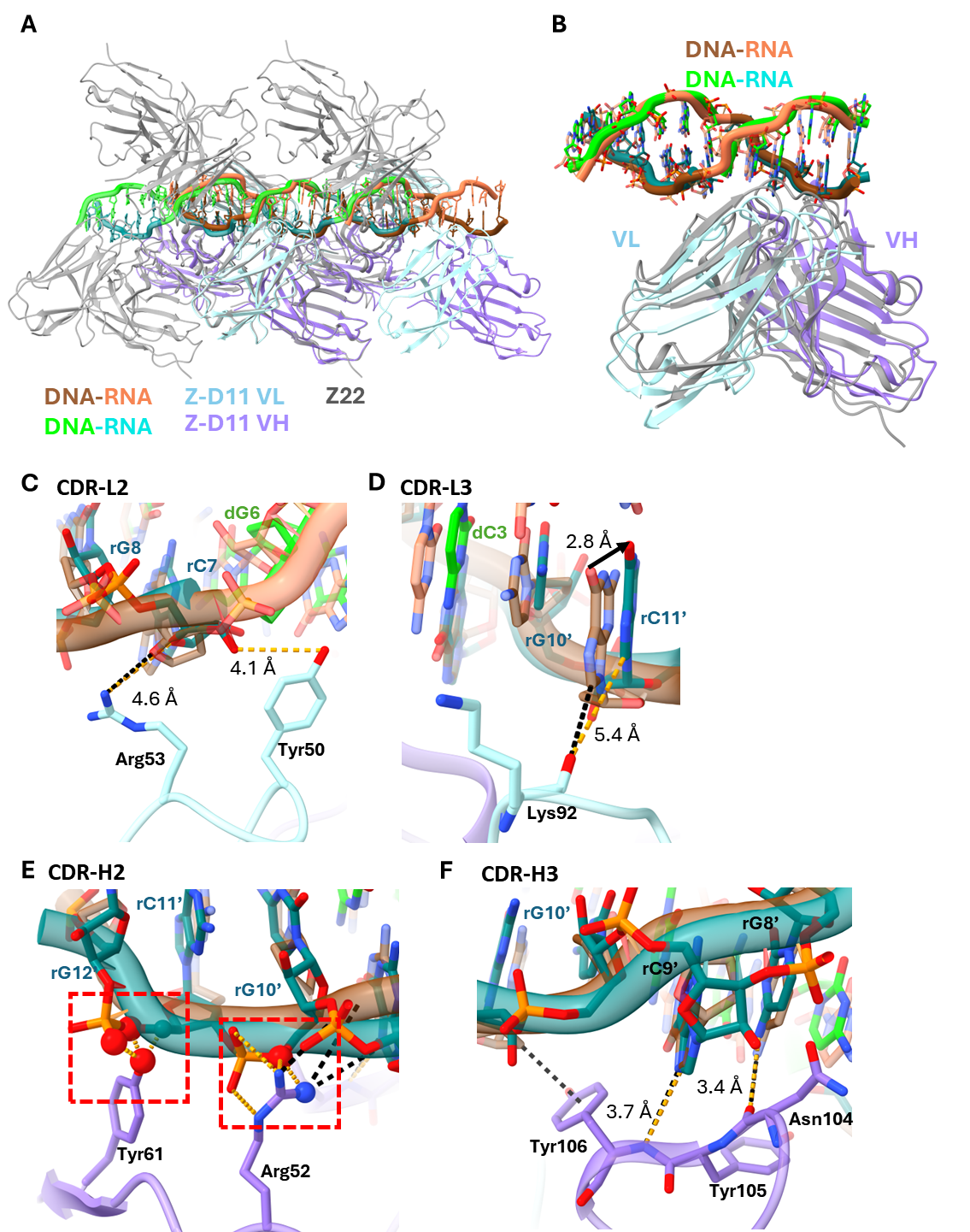

**Supplementary Figure S10. Optimization of Z-form induction conditions of d(CG)_6_ DNA and r(CG)_6_ RNA under physiological conditions.** Size exclusion chromatograms of Z-D11 in complex with r(CG)₆ under **A.** low salt Tris buffer (50 mM Tris pH 8, 150 mM NaCl); d(CG)₆ under **B.** low salt Tris buffer, and **C.** low salt HEPES buffer (20 mM HEPES pH 7.4, 150 mM NaCl). Size exclusion chromatograms of Z22 in complex with r(CG)₆ under **D.** low salt Tris buffer (50 mM Tris pH 8, 150 mM NaCl); d(CG)₆ under **E.** low salt Tris buffer, and **F.** low salt HEPES buffer (20 mM HEPES pH 7.4, 150 mM NaCl). The chromatograms for the complex, free antibody, and free NA are represented in green, blue, and yellow, respectively. Absorbance at 260 nm and 280 nm are presented as dashed and solid lines, respectively. Void volume for Superose 6 Increase 3.2/300 column is approximately 0.8 mL. Circular dichoism spectra of **G.** d(CG)₆ and **H.** r(CG)₆ in the presence of ADAR1 Zα under low salt buffer (20 mM HEPES pH 7.4, 150 mM NaCl).

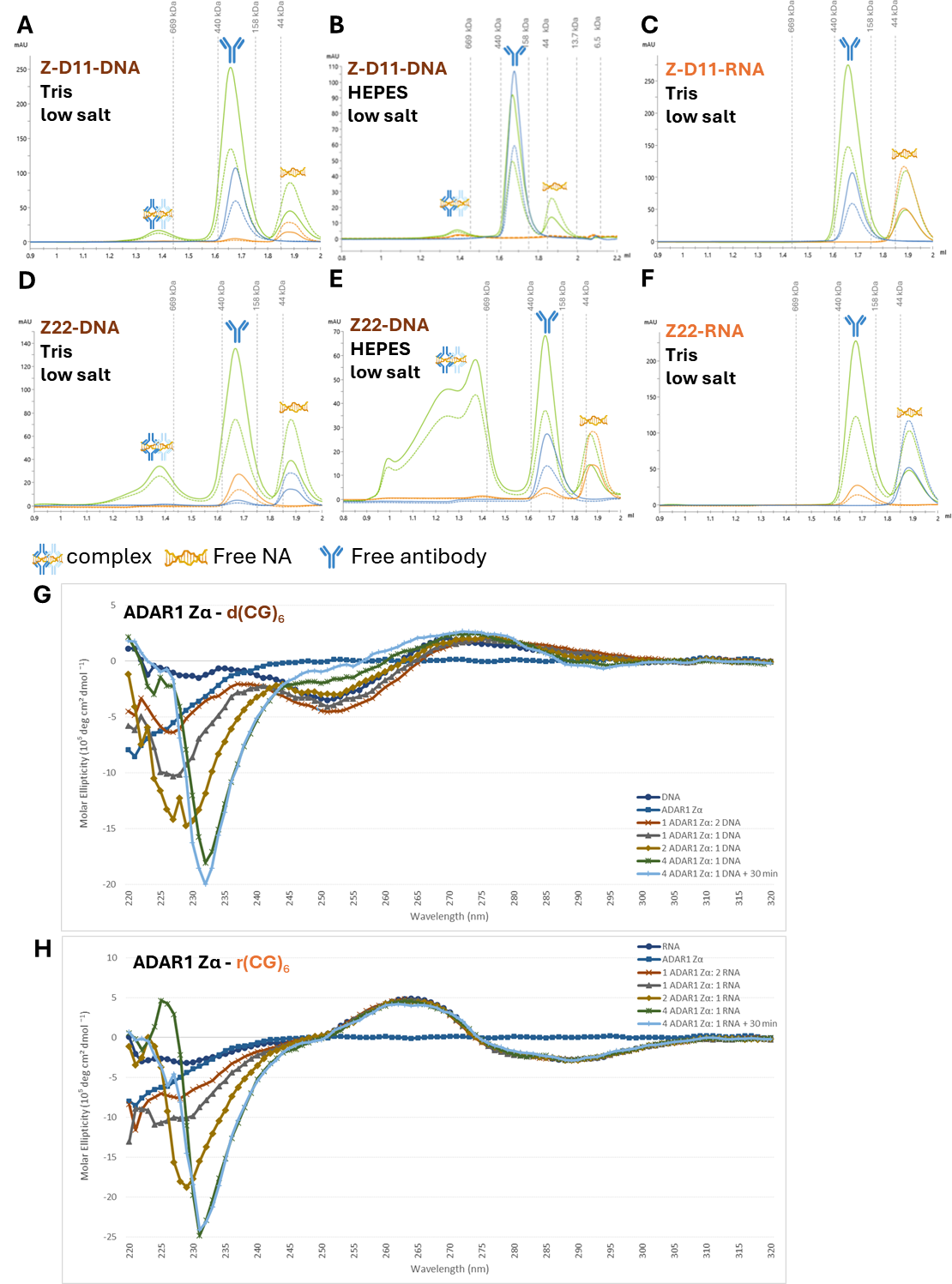

**Supplementary Table 1.** **Collection, refinement and validation statistics for cryo-EM density map and model.** Collection details for the cryo-EM map for Z-D11-drd(CG)_6_, Z-D11-rdr(CG)_4_, Z-D11-dr(CG)_6_, and Z22-rdr(CG)_4_ presented in the text are shown above, and validation statistics for the model generated from the map are shown below.

|  | **Z-D11-dr(CG)₆** | **Z-D11-rdr(CG)**_4_ | **Z22-dr(CG)₆** | **Z22-rdr(CG)**_4_ |
| --- | --- | --- | --- | --- |
| **EMDB:** | 81386 | 81320 | 81339 | 81340 |
| **PDB:** | 27SK | 27PG | 27QC | 27QD |
| **Data collection and processing** | | | | |
| Detector | Falcon 4i | | | |
| Magnification | 165,000 × | | | |
| Voltage (kV) | 300 | | | |
| Electron exposure (e–/Å^2^) | 50 | | | |
| Defocus range (µm) | -0.5 ~ -1.5 | | | |
| Pixel size (Å) | 0.76 | | | |
| Symmetry imposed | C1 | C1 | C1 | C1 |
| Initial particle images (no.) | 270,666 | 755,093 | 200,242 | 1,908,426 |
| Final particle images (no.) | 76,537 | 89,445 | 85,562 | 255,330 |
| Map resolution (Å) | 3.00 | 2.88 | 2.74 | 2.45 |
| FSC threshold micrographs | 0.143 | 0.143 | 0.143 | 0.143 |
| Map resolution range (Å) | 2.52 - 4.49 | 2.52 – 3.33 | 2.56 – 3.17 | 2.28 – 2.71 |
| **Refinement** | | | | |
| Initial model used (PDB code) | AlphaFold | AlphaFold | AlphaFold | AlphaFold |
| Model resolution (Å) | 2.0 | 2.0 | 2.0 | 2.8 |
| FSC threshold | 0.143 - 0.5 | | | |
| Model resolution range (Å) | 1.5 – 3.3 | 1.5 - 2.9 | 1.5 – 2.8 | 2.4-2.8 |
| Map sharpening B-factor (Å^2^) | 79.2 | 42.8 | 53.0 | 58.0 |
| **Model composition** | | | | |
| Non-hydrogen atoms | 8214 | 11942 | 11744 | 11757 |
| Protein residues | 916 | 1392 | 1373 | 1374 |
| Nucleotide | 48 | 48 | 48 | 48 |
| Water | 0 | 0 | 8 | 7 |
| Ligands | 0 | 0 | 0 | 0 |
| **B factors (Å^2^)** | | | | |
| Protein | 33.6/109.8/61.1 | 16.5/123.2/54.6 | 20.7/132.7/44.7 | 14.88/98.57/48.99 |
| Nucleotide | 43.4/133.2/65.2 | 2.2/60.2/24.2 | 19.3/45.8/27.1 | 14.95/106.78/35.14 |
| Water | - | - | 30.0/47.5/34.6 | 16.69/33.89/24.48 |
| Ligand | - | - | - | - |
| **R.M.S. Deviations** | | | | |
| Bonds length (Å) | 0.004 (0) | 0.012 (6) | 0.003 (0) | 0.002 (0) |
| Bonds Angle (˚) | 0.751 (0) | 1.054 (20) | 0.576 (0) | 0.675 (0) |
| **Validation** | | | | |
| MolProbity score | 2.15 | 1.60 | 1.31 | 1.15 |
| Clashscore | 6.61 | 6.73 | 3.93 | 3.13 |
| Rotamer outliers (%) | 2.66 | 2.01 | 0.68 | 0.51 |
| **Ramachandran Plot** | | | | |
| Favored (%) | 92.78 | 98.03 | 97.33 | 97.78 |
| Allowed (%) | 7.11 | 1.97 | 2.67 | 2.22 |
| Disallowed (%) | 0.11 | 0.00 | 0.00 | 0.00 |

**Supplementary Movie S1.** Spin video of the Z-D11-dr(CG)_6_ hybrid complex along the Z-NA axis. Z-DNA stretch is coloured in brown; Z-RNA stretch is coloured in coral. Z-D11 is coloured in purple and turquoise for the heavy chain and light chain, respectively.

(related to Figure 2B, 2C)

**Supplementary Movie S2.** Spin video of the Z22-dr(CG)_6_ hybrid along the Z-DNA axis. Z-DNA stretch is coloured in brown; Z-RNA stretch is coloured in coral. Z22 is coloured in silver and orange for the heavy chain and light chain, respectively.

(related to Figure 2E, 2F)

**Supplementary Movie S3.** Spin video of the Z-D11-dr(CG)₆ hybrid complex at the antibody-DNA interface. Z-DNA stretch is coloured in brown; Z-RNA stretch is coloured in coral. Z-D11 is coloured in purple and turquoise for the heavy chain and light chain, respectively.

(related to Figure 3D, Supplementary Figure S8A)

**Supplementary Movie S4.** Spin video of the Z22-dr(CG)₆ hybrid complex at the antibody-DNA interface. Z-DNA stretch is coloured in brown; Z-RNA stretch is coloured in coral. Z22 is coloured in silver and orange for the heavy chain and light chain, respectively.

(related to Figure 3E, Supplementary Figure S8B)

**Supplementary Movie S5.** Spin video of the Z-D11-rdr(CG)_4_ chimera complex along the Z-NA axis. Z-DNA stretch is coloured in brown; Z-RNA stretch is coloured in coral. Z-D11 is coloured in purple and turquoise for the heavy chain and light chain, respectively.

(related to Figure 4A, 4C)

**Supplementary Movie S6.** Spin video of the Z22-rdr(CG)_4_ chimera complex along the Z-NA axis. Z-DNA stretch is coloured in brown; Z-RNA stretch is coloured in coral. Z22 is coloured in silver and orange for the heavy chain and light chain, respectively.

(related to Figure 4B, 4D)

**Supplementary Movie S7.** Spin video of the Z-D11-rdr(CG)_4_ chimera complex at the antibody-DNA interface. Z-DNA stretch is coloured in brown; Z-RNA stretch is coloured in coral. Z-D11 is coloured in purple and turquoise for the heavy chain and light chain, respectively.

(related to Supplementary Figure S7C, S8C)

**Supplementary Movie S8.** Spin video of the Z22-rdr(CG)_4_ chimera complex at the antibody-DNA interface. Z-DNA stretch is coloured in brown; Z-RNA stretch is coloured in coral. Z22 is coloured in silver and orange for the heavy chain and light chain, respectively.

(related to Supplementary Figure S7D, S8D)

**Supplementary Movie S9.** Spin video of the modelled Z-D11-dr(CG)_6_ hybrid complex at the antibody-RNA interface. Black dashed lines represent the original interaction between Z-D11 and Z-DNA; Yellow dashed lines represent the predicted interaction between Z-D11 and Z-RNA. Z-DNA stretch is coloured in brown (original orientation) and lime green (rotated orientation); Z-RNA stretch is coloured in coral (original orientation) and teal (rotated orientation). Z-D11 is coloured in purple and turquoise for the heavy chain and light chain, respectively.

(related to Supplementary Figure S9)
